# The spliceosome assembles on excised linear introns to protect them from degradation

**DOI:** 10.64898/2026.01.21.700889

**Authors:** Glenn Wei Li, Max E. Wilkinson, David P. Bartel

**Affiliations:** Howard Hughes Medical Institute, Cambridge MA, USA; Department of Biology, Massachusetts Institute of Technology, Cambridge MA, USA; Whitehead Institute for Biomedical Research, Cambridge, MA, USA; McGovern Institute for Brain Research at MIT, Cambridge, MA, USA; Broad Institute of MIT and Harvard, Cambridge, MA, USA; Structural Biology Program, Memorial Sloan Kettering Cancer Center, New York, NY, USA

**Author notes:** Correspondence (M.E.W.), (D.P.B.).

## Abstract

In *Saccharomyces cerevisiae*, prolonged cellular stress induces some introns to accumulate post-splicing as stable, linear, spliceosome-protected RNAs^1^. These stable introns are defined by having short distances from their branchpoint (BP) sequences to their 3′-splice sites (3′SSs). Stable introns sequester splicing components, thereby reducing splicing activity and affecting cell growth in the stressed conditions. The mechanism by which these normally ephemeral products of pre-mRNA splicing persist cannot be explained by the current understanding of the splicing pathway, which derives primarily from studies of unstressed cells and their extracts^2,3^. Here, we determined the cryo-electron microscopy (cryo-EM) structure of a stable-intron complex purified from saturated-culture conditions. This structure and experimental follow-up show that a B^act^-like spliceosome protects stable introns from degradation, and that the short BP–3′SS distances of stable introns render this conformation of the spliceosome resistant to remodelling by helicases. Spliceosomes can also assemble onto artificial introns that have the same sequences as authentic stable introns but do not rely on splicing for their biogenesis, which demonstrates that spliceosomes arrive at this B^act^-like conformation by reassembling onto linear introns after their excision from pre-mRNAs. This reassembly activity is maintained in both stressed and unstressed cells. Thus, most yeast introns compete with pre-mRNAs for access to the splicing machinery, and budding yeast has co-opted this activity to adapt to environmental insults.

## Introduction

The spliceosome is a dynamic molecular machine that catalyzes the removal of introns from precursor mRNA (pre-mRNA). Each splicing event begins with the assembly of the spliceosome on the pre-mRNA and ends with dissociation of the spliced mRNA and disassembly of the intron-lariat spliceosome (ILS), with release of the lariat intron. This disassembly process liberates spliceosome components to re-assemble on a new pre-mRNA substrate and deprotects the lariat intron, which is then rapidly debranched and degraded^2,4^. In the budding yeast *S. cerevisiae*, prolonged cellular stresses, including those of saturated culture, induce a subset of introns to accumulate post-splicing as stable, linear RNAs (stable introns) that escape canonical debranching and decay^1^. Of the ∼300 introns of *S. cerevisiae*, approximately 30 accumulate as stable introns in saturated culture. These 30 introns, some of which accumulate to levels even higher than their host mRNAs, differ from the other introns in one key feature: They each have a short (typically <30-nt) distance between their branchpoint adenosine (BP) and 3′ splice-site (SS), a feature reported to be both necessary and sufficient to specify intron stability (Fig. 1a)^1^.

**Fig. 1:**
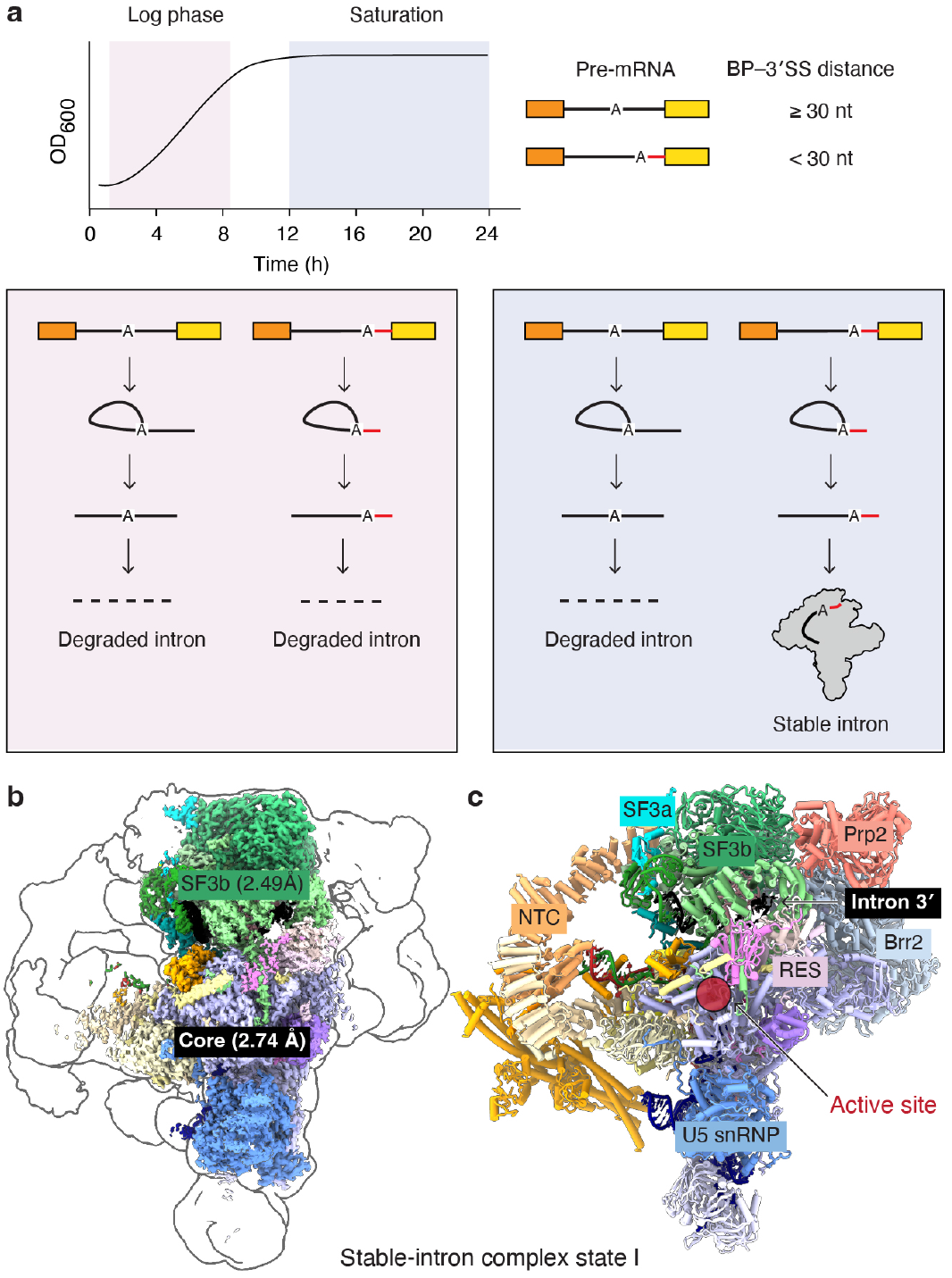
Cryo-EM structure of the stable-intron complex. **a**, Schematic of stable-intron accumulation. Excised introns with short BP–3′SS distances accumulate as spliceosome-protected linear species in saturated cultures. **b**, Composite density map for state I of the stable-intron complex. Shown are the focus-refined density maps of the spliceosome core and SF3b region (solid colours), and low-pass-filtered density for the core refinement (outline). **c**, Overall model for stable-intron complex state I.

In aggregate, stable-intron accumulation inhibits splicing and cell growth, which is thought to occur through titration of spliceosome components away from pre-mRNAs^1^. Indeed, stable introns co-sediment with complexes around 35– 50S, and stable introns purified from heavy-sedimenting fractions are bound by spliceosome components, including members of the Splicing Factor 3a (SF3a) and SF3b complexes, NineTeen Complex (NTC) and NineTeen Related complex (NTR)^1^. Based on the protein composition, the presence of the excised intron, and the absence of the spliced mRNA product in these complexes, stable introns are proposed to be bound by a ribonucleoprotein complex resembling the ILS, which protects them from the cellular RNA-decay machinery^1^. Thus, both the persistence and the function of stable introns depend on their prolonged association with spliceosome components.

How stable introns remain stably associated with the spliceosome post-splicing is unknown. Although slow ILS disassembly has been proposed to cause the accumulation of ILS complexes in some systems^5,6^, debranching in budding yeast is reported to occur only after ILS disassembly^7^. Thus, slowed ILS disassembly alone cannot account for the accumulation of debranched spliceosome-associated introns. In metazoan systems, however, debranching has been proposed to occur prior to spliceosome release through the recruitment of debranching enzyme (Dbr1) to the ILS by Aquarius (Aqr), a helicase that is absent in *S. cerevisiae*^8,9^. The stable-intron-spliceosome complex must also escape the disassembly activity of Prp43 and other DEAH-box helicases that can proofread spliceosomes assembled on aberrant splicing substrates^10^.

Hampering understanding of stable introns, which accumulate in saturated (not log-phase) cultures, is the lack of information regarding how the spliceosome structures and the yeast splicing pathway might change in saturated culture or in response to other long-term cellular stresses. All known structures of yeast spliceosomes have been generated from preparations of either endogenous splicing complexes purified from log-phase cultures or from splicing complexes generated invitro from extracts of log-phase yeast^11,12^. Moreover, most, if not all, studies involving biochemical dissection of the splicing pathway have been performed in log-phase yeast extracts^3^, a culture condition that does not support the formation of stable introns. To investigate the nature of the stable-intron complex and understand how it eludes the cellular mechanisms of spliceosome disassembly, we set out to use single-particle cryo-electron microscopy (cryo-EM) to determine the structure of a stable-intron complex purified from saturated culture.

## Results

### Stable introns reside in a B^act^-like spliceosome

We purified the stable-intron complex by expressing the *SAC6* stable intron in saturated culture and using an affinity tag on the intron to enrich for stable-intron complex and an affinity tag on the mRNA 5′ exon to deplete any un-spliced mRNAs (Extended Data Fig. 1a). The tagged *SAC6* intron accumulated as an excised linear intron in saturated cultures, similarly to the endogenous *SAC6* intron (Extended Data Fig. 1b). The affinity tag on the intron consisted of two MS2 hairpins, which enabled purification using the MS2-MBP fusion protein, and the affinity tag on the 5′ exon consisted of two PP7 aptamers enabling depletion of un-spliced mRNAs using Protein A-PP7 coat protein (Extended Data Fig. 1a). The purified spliceosomes contained fulllength excised linear introns, lacked un-spliced mRNAs, and contained snRNAs and proteins characteristic of active-site-containing spliceosomes (Extended Data Fig. 1c–f).

After collecting cryo-EM data from this sample (Extended Data Table 1), ab-initio reconstruction of a random subset of the 2 million picked particles yielded a volume with strong resemblance to the B^act^ spliceosome. Heterogeneous refinement of all particles using B^act^, C, and P-complex spliceosomes as references only yielded reconstructions in a B^act^ conformation (Extended Data Figs. 2 and 3). Many aspects of the reconstruction confirm that the stable-intron complex is B^act^-like (Fig. 1b and Extended Data Fig. 4a,b): The stable-intron complex contained the U2 snRNP (including the SF3a and SF3b complexes), U5 snRNP, U6 snRNA, the NTC and NTR complexes, the Retention and Splicing (RES) complex, intron RNA, and several B^act^-specific splicing factors, notably Cwc24 and the DEAH-box helicase Prp2, which is positioned near the intron 3′ end (Fig. 1c). Moreover, the intronic portion of the 5′SS is held in the RNA active site, whereas the branchpoint is held ∼50Å away from the active site, shielded by the SF3b complex.

Alongside these features all shared with yeast and human B^act^ spliceosomes^13–16^, the stable-intron complex differed from the canonical B^act^ complex in two important ways—one at the 5′SS and the other at the U6 snRNA internal stem-loop (ISL)—both of which are described in next section. Some minor differences with canonical B^act^ were also observed (Extended Data Fig. 4a). Our refinement yielded two distinct states of the B^act^-like stable-intron spliceosome, both with similar particle numbers (Extended Data Fig. 2). State I was most like canonical B^act^, whereas state II, had further differences with canonical B^act^ (Extended Data Fig. 4a), in that it lacks Brr2 (Extended Data Fig. 4a,b), has more flexibility of the SF3b complex relative to the rest of the spliceosome (Extended Data Fig. 4c), and has a shift in Prp8 endonuclease domain that causes this domain to more closely resemble its position in the C complex or P complex (Extended Data Fig. 4d)^11,17–19^. In both states I and II, a loop in the Prp8 endonuclease domain (residues 1615 – 1624) is remodelled such that Tyr1620 can stack on the RNA bound to the U5 loop I, resembling its conformation in C or P complexes (Extended Data Fig. 4e). Lastly, whereas the B^act^ complex and state I of the stable-intron complex contain Cwc24, a protein factor that recognizes and stabilizes the GU at the 5′SS^20^, state II lacks density for Cwc24 and the intron 5′ end (Extended Data Fig. 4f).

Despite these differences, both states of the stable-intron complex were strikingly similar to the B^act^ spliceosome, which was surprising, as the stable-intron complex was expected to resemble a spliceosome bound to an excised intron lariat (i.e., an ILS) and not a conformation that is normally assembled upon un-spliced pre-mRNAs.

### The stable-intron complex differs from the B^act^ spliceosome at the 5′SS and active site

In the B^act^ spliceosome, the 5′ exon and 5′SS are held in the RNA active site prior to cleavage and branch formation, with the 5′ exon bound to loop I of U5 snRNA and the 5′SS bound to the ACAGAGA box of U6 snRNA^14–16^. In the stable-intron complex, we still observed cryo-EM density for an RNA basepaired to U5 snRNA loop I, and this density was particularly strong in state II, accompanied by stronger density for the splicing factor Cwc21, which lines the exon-binding channel (Extended Data Fig. 4a). However, several lines of evidence showed that this density did not correspond to 5′ exon linked to the intron prior to cleavage. First, primer-extension analysis indicated that the purified stable-intron complex contained only full-length excised intron, with no evidence of 5′ exon appended to the intron 5′ terminus (Extended Data Fig. 1d). Second, in both states I and II, we observed a clear gap in cryo-EM density between the intron 5′ phosphate and the 3′ OH of the RNA bound to the U5 loop I (Fig. 2a,b). Third, the 2.6 Å resolution cryo-EM map of state II unambiguously discriminated between purines and pyrimidines at the 3′-terminal sequence of the RNA bound in the exon-binding channel, and these differed from the sequence of the 5′ exon of our expression construct (Extended Data Fig. 5a). Instead, they matched the 3′-end sequence of *CUT314*, an oligo-adenylated non-coding RNA that was abundant in our purifications of stable-intron complex (Extended Data Fig. 5b,c). However, binding of *CUT314* did not appear to be an important part of the stable-intron pathway, in that deleting *CUT314* had no effect on stable-intron accumulation (Extended Data Fig 5d), which suggested that other 3′-terminal polyadenylated molecules might substitute for *CUT314*. In sum, our results indicated that for both state I and II the structure corresponded to a linear excised intron bound to a B^act^-like spliceosome, together with another RNA fragment mimicking the 5′ exon.

**Fig. 2:**
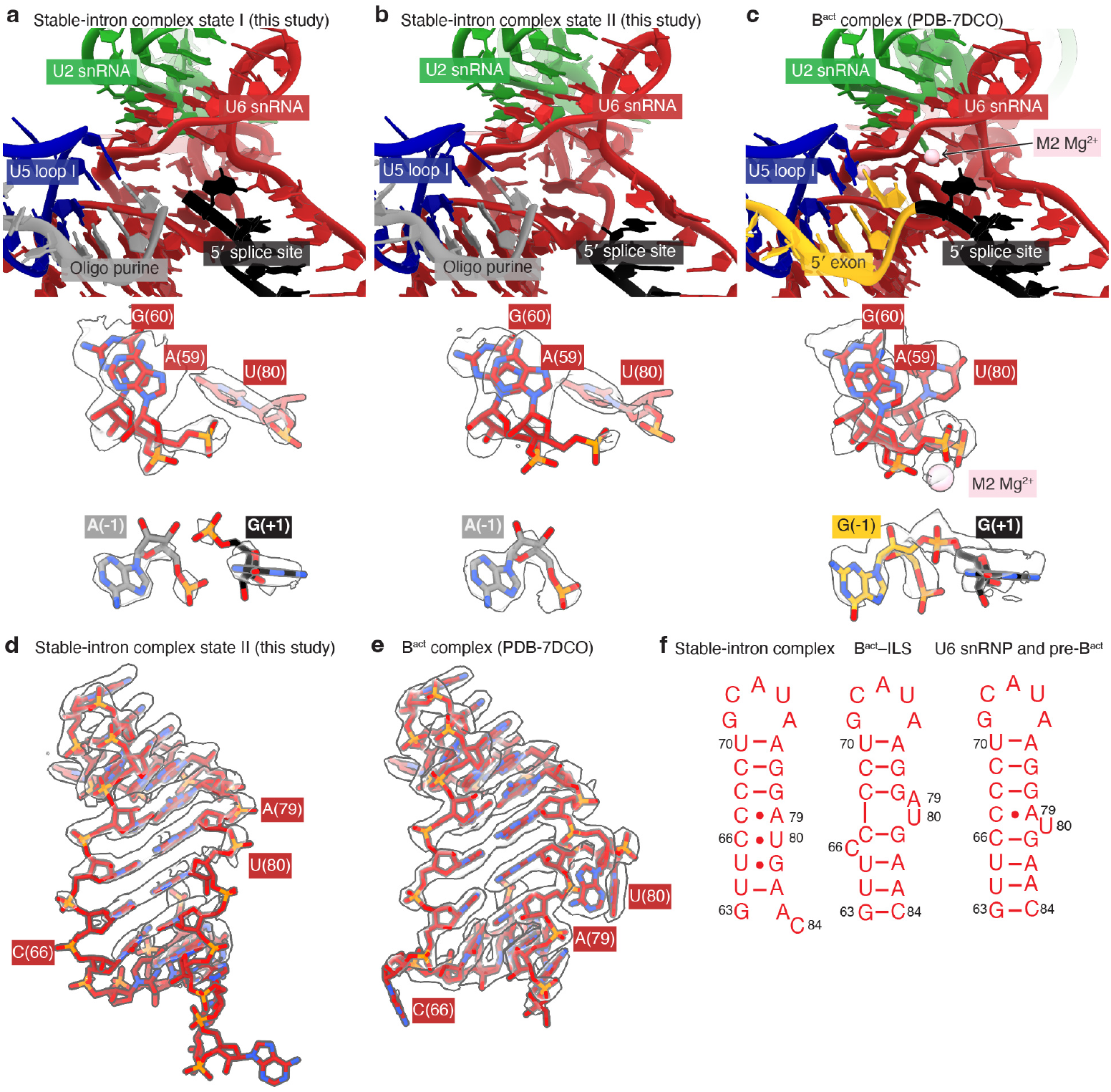
Stable-intron complex contains a remodeled active site. **a**, Active-site organization for state I of the stable-intron complex. Shown are the overall model for the RNA active site (above) and the cryo-EM density for the active site residues (below). **b**, Active-site organization for state II of the stable-intron complex; otherwise, as in **a. c**, Active-site organization of the yeast B^act^ complex in a view analogous to that of **a. d**,**e**, Structure of the U6 internal stem loop (ISL) in either the stable-intron complex (**d**) or the B^act^ complex (**e**) in their respective cryo-EM densities. **f**, Secondary structure of the U6 ISL in the indicated spliceosome structures^2,22–24^. Dots indicate non-Watson–Crick base pairs.

Accompanying these differences at the 5′SS were key differences at the active site. During the normal splicing cycle, the RNA active site is formed during the transition from the B to the B^act^ complex and persists until disassembly of the ILS at the end of the cycle^2^. The U6 internal stem-loop (ISL), an element that also persists from B^act^ to ILS, is critical for formation of the active site; it places nucleotide U80 in position to coordinate the M2 magnesium ion, which stabilizes growing charge on the scissile phosphate at the exon–intron boundary during nucleophilic attack (Fig. 2c)^21^. In the stable-intron complex, the U6 ISL and the RNA active site adopt alternative conformations, with loss of the M2 magnesium (Fig. 2a–c). The A79 and U80 bases, which are normally flipped out of the ISL, instead form mismatches with C66 and C67, and the register of the pairing in the remainder of the stem is adjusted to accommodate this change (Fig. 2d– f). These changes to the RNA active-site structure presumably render the stable-intron complex catalytically inactive.

### The structure near the intron 3′ terminus suggests a role for Prp2 access

We were particularly interested in the structure of the stable-intron complex in the vicinity of the intron 3′ terminus, as having a short (typically < 30-nt) distance between the BP and 3′ SS licenses introns to become stabilized in saturated culture and certain other stresses^1^. With the important exception of the missing 3′ exon, the overall architecture of the stable-intron complex surrounding the intron 3′ terminus was indistinguishable from that of the B^act^-complex (Fig. 3a). As in the B^act^-complex, the SF3b complex binds the intron BP and the downstream 15 nucleotides, and RES-complex protein Ist3 binds to the five terminal nucleotides of the *SAC6* intron (nucleotides 15–19 downstream of the BP), in an interaction stabilized by contacts with Bud13, Prp45 and Prp8^15,16,25,26^. Also as the in B^act^-complex, Prp2, a DEAH-box helicase, is positioned nearby, through interactions with the SF3b-complex components Rse1 and Hsh155^14–16^. In the normal B^act^ complex, the pre-mRNA continues into the active site of Prp2, and Prp2 translocates 3′–5′ along the pre-mRNA, which destabilizes the branch helix, SF3a, SF3b and RES complexes, thereby inducing the transition from B^act^ to B*^22,27^. Biochemical experiments indicate that Prp2 binds the pre-mRNA beginning 20–28-nt downstream of the BP^28,29^ and requires a BP–3′-end distance of at least 25–30 nucleotides to efficiently catalyze the transition to B*^28,30–32^, a range overlapping with that inferred by structures showing that Prp2 binds the pre-mRNA 29–32 nucleotides downstream of the BP^14^.

**Fig. 3:**
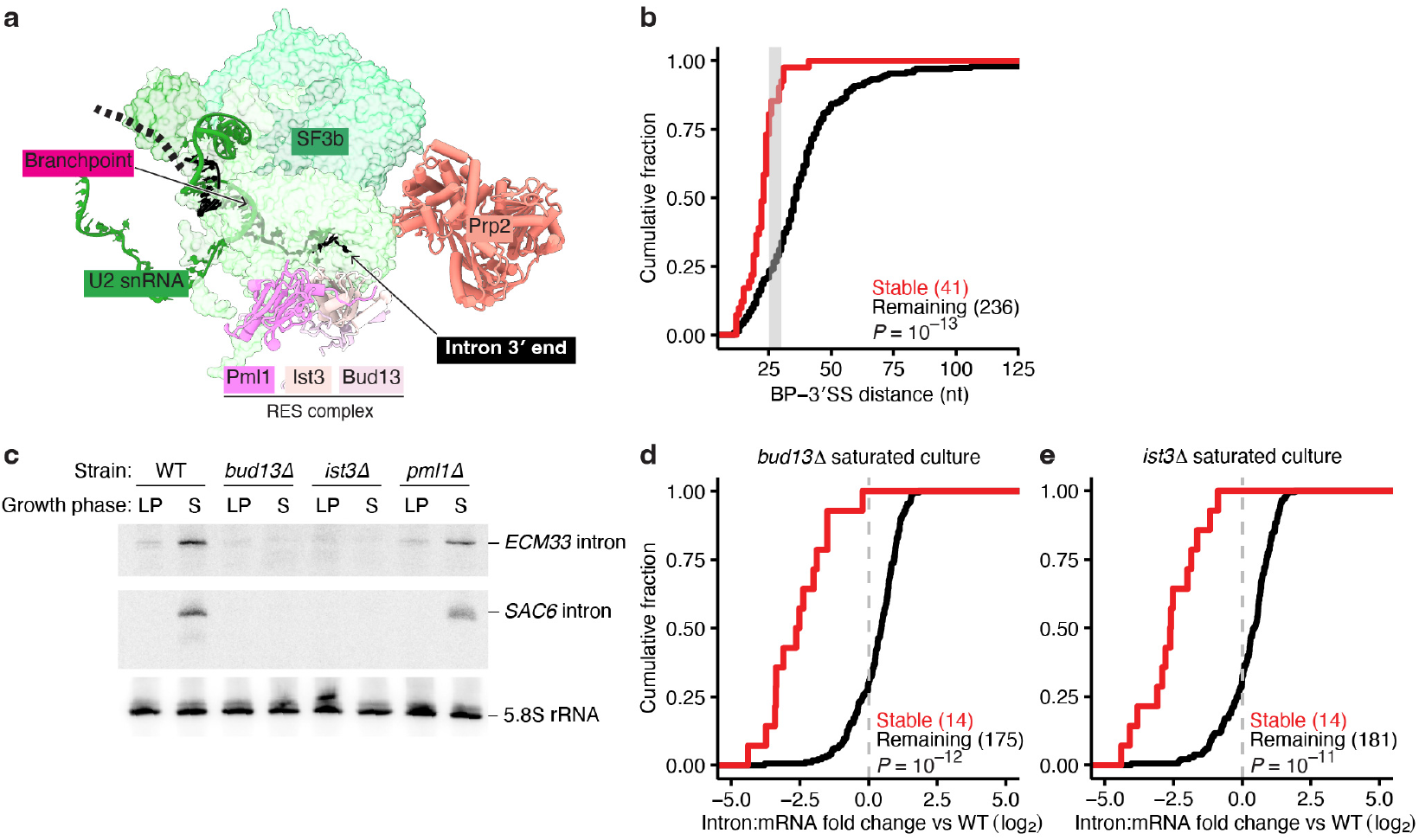
The RES complex protects the stable-intron complex from remodelling by Prp2. **a**, Structure of the stable-intron complex near the intron 3′ terminus. Factors not in the immediate vicinity of the intron RNA are hidden for clarity. **b**, Tendency of stable introns to have BP–3′SS distances too short to enable efficient Prp2-mediated remodeling. Plotted are cumulative distributions of BP–3′SS distances for stable introns and the remaining introns (*P* value, two-tailed Kolmogorov–Smimov test). The 41 stable introns were identified either previously^1^ or in the current study (Extended Data Table 2). The gray rectangle denotes range of minimum BP–3′-end distances reported for the Prp2-dependent branching reaction^30,32^. **c**, Reduced accumulation of *ECM33* and *SAC6* stable introns in RES-complex deletion mutants. Shown is an RNA blot of a denaturing gel resolving total RNA from log-phase (LP) and saturated (S) cultures of wildtype (WT) and RES-complex mutant strains. The blot was probed for the *ECM33* (top) and *SAC6* (middle) introns simultaneously and then re-probed for the 5.8S rRNA (bottom) as a loading control. Blot is representative of *n* = 3 biological replicates. **d**,**e**, Global reduction of stable-intron accumulation in *bud13Δ* (**d**) and *ist3Δ* (**e**) strains compared to the wildtype strain in saturated culture. Plotted for both the 14 stable introns and the remaining introns are cumulative distributions of fold changes of the intron:mRNA ratio observed between saturated cultures of the mutant and WT strains. Results are shown for introns for which both the intron and the host mRNA met a 10-read cutoff in at least half of the sequencing libraries analyzed. For each fold change, the mean of two biological replicates is plotted (*P* values, two-tailed Kolmo-gorov–Smimov test). The stable introns were the 14 identified in the matched wildtype control strain (Extended Data Table 2). The higher intron:mRNA ratios for most non-stable introns in the mutant strains are indicative of increased intron retention in these strains.

In the stable-intron complex, Prp2 is bound in its normal position but cannot access the 3′ end of the stable intron, as the *SAC6* intron, with its short BP–3′SS distance, terminates within the RES complex (Fig. 3a). The length requirement for stable-intron accumulation aligns well with the distance requirement for Prp2 function: of the 41 stable introns identified over a range of stresses^1^ (Extended Data Table 2), all but four have BP–3′SS distances of 29 nucleotides or less (Fig. 3b). This short distance would make most stable introns bound in a B^act^-like state poor substrates of Prp2-mediated remodeling. Conversely, non-stable introns, by virtue of their longer BP–3′SS distances, would remain accessible to Prp2, even without a 3′-exon. Thus, when considering our structure together with previous structural and biochemical findings, we suggest that stable introns accumulate in a B^act^-like state because their BP–3′SS distances are too short to permit translocation by Prp2, and this differential accessibility to Prp2-catalyzed remodeling enables the spliceosome to discriminate between stable and non-stable introns.

### The RES complex is necessary for stable-intron accumulation

A structure of the human spliceosome stalled between B^act^ and B* indicates that Prp2 binds to branchpoint-proximal nucleotides as it translocates along the intron, which displaces the RES complex from the intron 3′-end^22^. The RES complex is composed of Bud13 and Ist3, which stabilize the intron 3′-end^33^, and Pml1, which has a more minor role in splicing and instead is involved in the retention of unspliced mRNAs in the nucleus^34–36^. Our model of stable-intron accumulation predicts that without the RES complex, Prp2 could access the stable intron 3′-end, remodel the stable-intron complex, and thereby facilitate the release of the intron from the protective association with the spliceosome. To test this prediction, we examined the accumulation of two stable introns (*ECM33* and *SAC6*) after deleting components of the RES complex. Deleting either *BUD13* or *IST3* substantially reduced accumulation of the *ECM33* and *SAC6* stable introns in saturated cultures, whereas deleting *PML1* had little effect, consistent with Pml1 having a smaller role in directly regulating splicing (Fig. 3c).

To extend this analysis to the other stable introns that accumulate in saturated culture, we performed RNA-seq on *bud13Δ, ist3Δ*, and wild-type control strains in log-phase and saturated cultures. Using our bioinformatic pipeline for identifying stable introns from RNA-seq data, we detected 14 stable introns from analysis of the wild-type control cultures but no stable introns from analysis of strains with deletions in *bud13* and *ist3*, as expected if these components of the RES complex were required for stable-intron accumulation (Extended Data Table 2). Moreover, when comparing saturated cultures from the different strains, we observed a global loss of stable-intron accumulation in mutants of the RES complex compared to the wild-type strain (Fig 3d,e). This change occurred without a corresponding change in the expression of exons that flank stable introns, indicating that the RES complex components are necessary for the accumulation of stable introns but do not dramatically affect the splicing of pre-mRNAs with stable introns (Extended Data Fig. 6a). Upon loss of *BUD13* and *IST3*, the absolute accumulation of stable introns was indistinguishable from that of other introns in saturation, which indicated that the large decrease in stable-intron accumulation as compared to other introns was not solely due to the high expression of stable introns in wild type cells as compared to other introns (Extended Data Fig. 6b). The effects on intron accumulation were specific to stable introns and to saturated cultures, i.e., in log-phase, intron accumulation changes observed upon knocking out RES-complex components did not differ between stable intron and other introns (Extended Data Fig. 6c), and after knocking out RES-complex components, accumulation of non-stable introns did not decrease in both log-phase and saturated cultures (Fig. 3d,e and Extended Data Fig. 6c). Instead, we observed an increase in non-stable intron accumulation (Fig 3d,e and Extended Data Fig. 6c), which can be attributed to decreased efficiency of splicing in RES-complex mutants causing increased intron retention, as reported previously^35,37^. We conclude that both *BUD13* and *IST3* are necessary for stable-intron accumulation, which supports our model that they protect the 3′-end of stable-introns from the activity of Prp2.

### Mechanistic possibilities for how stable introns arrive at a B^act^-like state

The structure of the stable-intron complex explained the protection of stable introns: they are sequestered in a B^act^-like complex recalcitrant to Prp2-mediated remodeling due to a short BP–3′SS distance. However, this explanation raised the even more perplexing question of how the spliceosome arrives at this off-pathway conformation assembled on excised, debranched introns. We envisioned two possible mechanisms. In the first, a spliceosome associated with the excised lariat intron (i.e., the ILS) somehow permits debranching and then undergoes a series of reverse-splicing-like steps to transition back to a B^act^-like state, but cannot proceed further back beyond this state due to the challenge in reversing the extensive rearrangements that occur during the B-to-B^act^ transition, especially the Brr2-mediated removal of the U4 snRNA^38^. The feasibility of this mechanism is supported by the observation that both catalytic steps of splicing are reversible in vitro^39^ and that the two catalytic states of the spliceosome exist in equilibrium^40,41^. However, reversion from B* to B^act^ has not been observed and is expected to be particularly challenging due to the large-scale rearrangements of the BP and SF3b complex during this transition^2^.

In the second mechanism, the lariat precursor of the stable intron dissociates from the spliceosome, is debranched, and then the excised linear intron associates with a new spliceosome through a pathway resembling canonical spliceosome assembly. In-vitro spliceosome assembly requires neither a 5′ exon^42^ nor a 3′ exon^30,32^. Thus, in principle, an excised linear intron that lacks both exons could enter the spliceosome pathway and progress until insufficient RNA for Prp2-mediated remodeling traps the complex in the B^act^-like state. This mechanism implies that, in some conditions, spliceosome reassembly can at least to some degree compete with intron degradation, perhaps through protection of the free intron from cellular nuclease activity.

### Stable-intron accumulation depends on U1 snRNP proteins

Canonical spliceosome assembly begins with the recognition of the 5′SS by the U1 snRNP, recognition of the branch sequence by the Mud2-Msl5 heterodimer, and recognition of the 5′ 7-methyl-GTP cap structure to form the E complex^43– 45^. Genetic and biochemical data indicate that the E-complex components compensate for each other when faced with a suboptimal substrate or when one or more of the components are mutated. For example, several components of the yeast U1 snRNP are not essential for viability or for splicing canonical pre-mRNA substrates, but are required for splicing suboptimal substrates, including pre-mRNAs that contain mismatches to the well-defined yeast 5′SS and branch site (BS) sequences, and pre-mRNAs that lack a 5′ 7-methyl-GTP cap^46–51^. We hypothesized that if stable introns access the B^act^-like complex through reassembly, linear excised introns, which contain intact splice-site sequences but lack exons and a 5′ cap structure, would behave like suboptimal splicing substrates and thus would depend on non-essential components of the U1 snRNP to re-associate with the spliceosome.

To test this prediction of the reassembly model, we examined the accumulation of stable introns in four strains with defective splicing of pre-mRNAs with suboptimal splice sites: *snu56-2*, a hypomorphic allele of the essential yeast-specific U1 snRNP component *SNU56*^46^, *mud1Δ*, a deletion mutant of the yeast U1-A homolog *MUD1*^52^ and *nam8Δ*, a deletion mutant of the non-essential *NAM8* gene required for the splicing of meiotic transcripts^48,53^, as well as *mud2*Δ, a deletion mutant of the E-complex protein *MUD2*, the yeast U2AF65 homolog, which helps recognize the BS^49^. All tested U1 snRNP mutant strains had decreased accumulation of both *ECM33* and *SAC6* stable introns (Fig. 4a,b), which indicated that robust 5′SS recognition was necessary for the accumulation of stable introns. *MUD2* deletion resulted in decreased *SAC6* but not *ECM33* intron accumulation, which suggested that BS recognition might be relatively less important than 5′SS recognition in the reassembly of excised introns, and was in line with the observation that perturbations to the spliceosome can result in intron-specific changes in expression^54^.

**Fig. 4:**
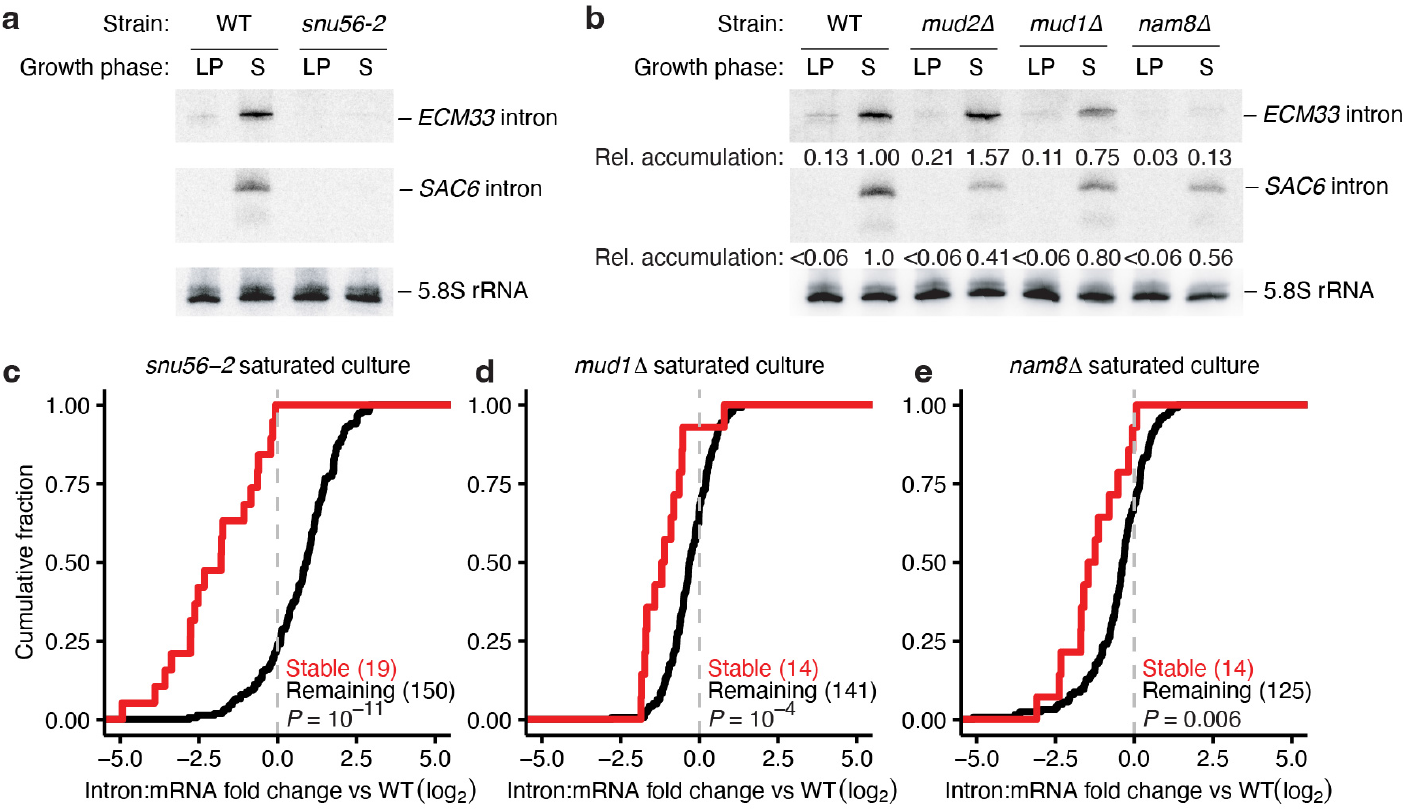
Stable-intron accumulation depends on U1 snRNP proteins. **a**, Reduced accumulation of *ECM33* and *SAC6* stable introns in the *snu56-2* mutant strain; otherwise as in Fig. 3c. *n* = 3 biological replicates. **b**, Reduced accumulation of *ECM33* and *SAC6* stable introns in mutants of the E-complex and U1-snRNP; otherwise as in (A). Numbers below the *ECM33* and *SAC6* blots indicate the mean level of the stable intron normalized to the WT saturated-culture condition after first normalizing all lanes to the 5.8S rRNA loading control. *n* = 3 biological replicates. **c**–**e**, Global reduction of stable-intron accumulation in *snu56-2* **(c)**, *mud1Δ* (**d**) and *nam8Δ* (**e**) strains compared to wildtype; otherwise as in Fig. 3d.

To examine the transcriptome-wide effects of these perturbations on stable-intron accumulation, we performed RNA-seq on the U1 snRNP mutants in both log phase and saturated cultures. Consistent with the U1 snRNP being necessary for the accumulation of stable introns, we identified fewer stable introns in the mutant strains than in the matched wildtype controls; we identified no stable introns in *snu56-2* compared to 19 in its matched wildtype control and six stable introns in the *mud1Δ* strain and three stable introns in the *nam8*Δ strains as compared to 14 in their shared wildtype control sample (Extended Data Table 2). For all three strains examined, accumulation of stable-introns decreased transcriptome-wide compared to the wildtype strain (Fig. 4c–e) and was not accompanied by dramatic changes in the levels of the flanking exons (Extended Data Fig. 7a), indicating that the U1 snRNP proteins examined are required for the accumulation of stable introns but not for splicing of the pre-mRNAs from which stable introns derived. Moreover, as with the RES-complex mutants, the effect of U1 snRNP mutation was specific to stable introns and to saturated cultures (Fig. 4c–e, Extended Data Figure 7b,c).

In both the RNA blot and RNA-seq analyses, the effect of the *snu56-2* mutation on stable-intron accumulation was more severe than that of deleting *NAM8* or *MUD1*. One possible explanation is that the hypomorphic *snu56-2* caused a greater perturbation to U1 snRNP function. This interpretation is supported by the reduced splicing efficiency in the *snu56-2* mutant, as evidenced by an increased intron-to-exon ratio for most non-stable introns (Fig. 4c)—a pattern diagnostic of intron retention not observed in the *mud1*Δ and *nam8*Δ mutants, which instead show a mild decrease in non-stable intron abundance (Fig. 4d,e). *MUD1* and *NAM8* are not required for vegetative splicing^50,52^, and the decrease in non-stable intron abundance could reflect increase in overall splicing efficiency due to the decreased spliceosome sequestration by stable introns.

In sum, the dependence of stable-intron accumulation on U1 snRNP proteins that are not required for the splicing of canonical pre-mRNA substrates supports the reassembly model and not the reverse-splicing model, in that reassembly but not reverse splicing would pass through U1 snRNP-facilitated formation of the E complex. These results further suggest that U1 snRNP must recognize a suboptimal, exon-free 5′SS to direct spliceosome assembly onto the excised intron and that a compromised U1 snRNP is unable to effectively act on these suboptimal substrates.

### Stable-intron accumulation depends on 5′SS and branch site strength

The U1 snRNP recognizes the 5′SS through base-pairing interactions with the U1 snRNA^44^. U1 snRNP recruitment to the 5′SS is enhanced in the presence of a 5′ cap structure and BS by the other E-complex proteins, and in yeast, this enhancement is particularly important for stable U1 binding when the 5′SS is mutated away from the well-conserved consensus sequence^51^. We reasoned that if stable introns accessed the B^act^-like assembly through the spliceosome-reassembly mechanism, then these excised introns, which lack 5′ caps, would be especially sensitive to suboptimal 5′SS and BS sequences. Examination of the splice site sequences of the 41 stable introns identified to date revealed that stable introns were enriched for having consensus 5′SS and BS sequences both compared to all introns and compared to introns with short BP–3′SS distances (Fig. 5a, Extended Data Figure 8a,b). Although low transcription during the time permissive for stable-intron accumulation can explain why certain introns with short BP–3′SS distances are not identified as stable^1^, this analysis indicated that most introns that fail to accumulate despite short BP–3′SS distances are not stabilized due to suboptimal branch site or 5′SS sequences.

**Fig. 5:**
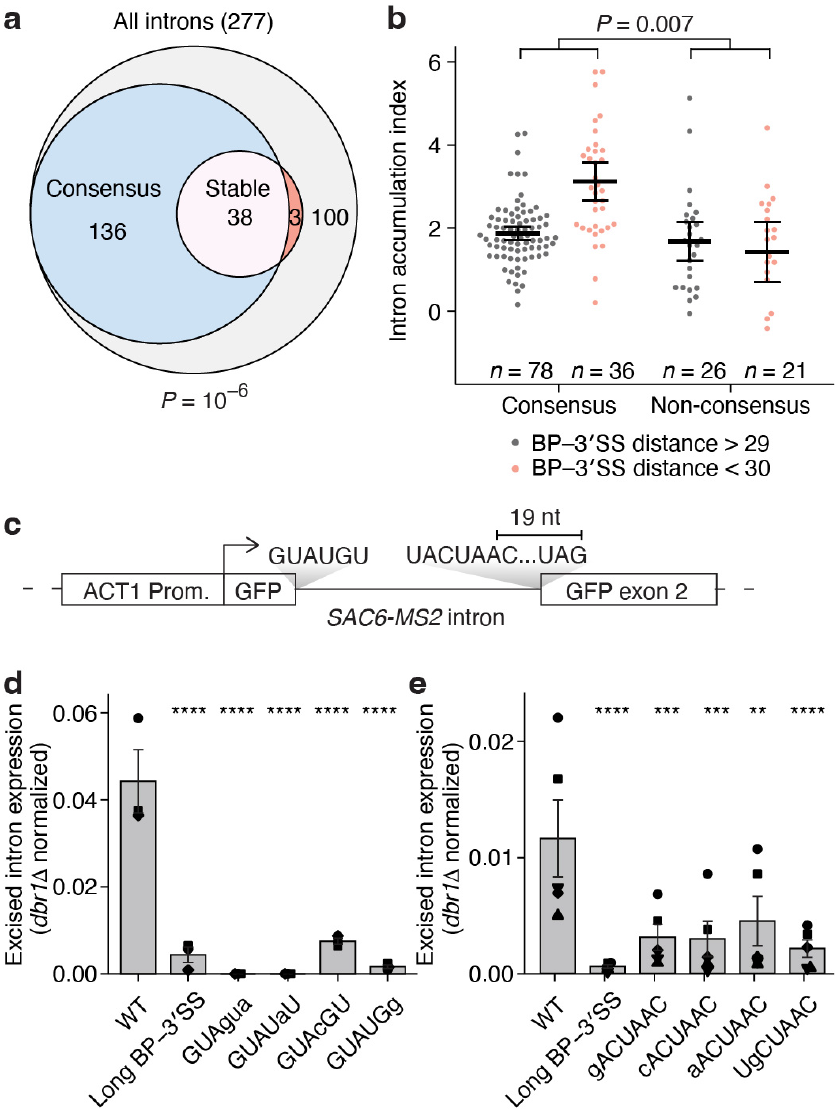
Consensus 5′ splice-site and branch-site sequences sub-stantially enhance stable-intron accumulation. **a**, Enrichment of consensus 5′SS and branch site sequences in stable introns. Venn diagram shows overlap between stable introns and introns containing consensus 5′SS and branch site sequences. (*P* = 10^−6^, hypergeometric test). **b**, Lack of preferential accumulation of introns with short BP–3′SS distances but non-consensus 5′SS or branch site sequences. Plotted are mean intron accumulation indices (*n* = 6 biological replicates), which were calculated from the fold change in intron accumulation observed between saturated and log-phase cultures, normalized by the fold change in mRNA accumulation. Results are shown for introns for which both the intron and the host mRNA met a 10-read cutoff in at least half of the sequencing libraries analyzed. Black bars denote the mean; error bars denote the 95% confidence interval. The number of introns in each category is noted below each set of points. The point for the *SNR17B* intron (which has a non-consensus branch site sequence, a BP–3′SS distance of 13, and an intron accumulation index of –3.66) fell outside the axis. *P* value, two-tailed t-test comparing the difference between short and long BP–3′SS introns for introns with consensus versus non-consensus splice sites, difference between groups, 1.4, 95% confidence interval, 0.41 to 2.5. **c**, Reporter intron design. The *ACT1* promoter was used to drive expression of a *GFP* mRNA that contained the MS2-tagged *SAC6* intron. **d**,**e**, Reduced stable-intron accumulation upon mutation of the 5′SS (**d**) and the branch site (**e**). Plotted for each variant is the mean abundance in saturated culture, as a fraction of the abundance in *dbr1Δ* (which indicates intron production), after normalizing to the signal for 5.8S rRNA (a loading control). Error bars, s.e.m. Shapes represent each of three (**d**) or five (**e**) biological replicates. **, *P* < 10^−2^; ***, *P* < 10^−3^; ****, *P* < 10^−4^, one-way paired ANOVA (equal variances assumed) with Dunnett’s multiple comparisons tests comparing each variant to wildtype. See Extended Data Fig. 8c,e for representative blots.

To examine further the effects of these sequence features on endogenous introns, we re-analyzed the RNA-seq datasets from the wildtype strain, with a focus on the intron accumulation index, which quantifies the change to the intron-to-mRNA ratio observed between saturated and logphase cultures. Although this measure of intron accumulation was one component of our pipeline for identifying stable introns, it did not solely determine whether an intron was classified as stable or not. We grouped introns by whether they possessed a short BP–3′SS distance and whether they contained consensus 5′SS and BS sequences and evaluated whether these sequence features associated with increased intron accumulation (Fig. 5b). Among introns with consensus 5′SS and BS sequences, those with short BP–3′SS distances typically underwent greater saturation-dependent increases in abundance than did their long BP–3′SS counterparts, consistent with the known effect of BP–3′SS distance on intron accumulation in saturated cultures^1^. In contrast, introns that lacked a consensus 5′SS or BS sequence did not show this difference in accumulation between long and short BP–3′SS distances. These results refuted our previous conclusion that a short BP–3′SS distance is sufficient for stabilization of an excised intron in saturated culture^1^, in that this stabilization also requires strong splicesite sequences.

To directly assay the effects of 5′SS and BS sequence on stable-intron accumulation, we designed a series of reporter introns that were based on the MS2-tagged *SAC6* intron but contained non-consensus splice-site sequences chosen either from the introns of meiotic pre-mRNAs that depend on *NAM8* for their splicing^48^ or from introns with short BP–3′SS distances with low intron accumulation indices (Fig. 5b). We also included a variant of the *SAC6* intron with a lengthened BP–3′SS distance (34 compared to 20-nt) to compare the effects of SS sequence mutation to the previously reported effect of BP–3′SS distance mutation^1^. For all tested variants, we controlled for potential changes in splicing efficiency, which affects stable-intron accumulation for the simple reason that introns must be excised before they can be stabilized. Accordingly, changes in linear excised introns were normalized to changes in the accumulation of the lariat intron in the *dbr1Δ* strain, which, due to the lack of intron debranching and decay^55^, should depend on the relative efficiency of excision by the spliceosome, or on any change in production of the mature mRNA from which the intron was spliced.

The reporter intron accumulated as a linear excised intron in saturated cultures, and lengthening the BP–3′SS distance disrupted accumulation of the full-length intron (Fig. 5d,e, Extended Data Fig. 8c,e). Not disrupted was accumulation of a lower-molecular-weight product that appears to correspond to an intron species that was trimmed from the 3′ end, in that it did not bind to a probe that hybridizes to the extreme 3′ end of both the wild-type and mutant introns (Extended Data Fig. 8c,e). This hinted that processing of the intron 3′ end by the nuclear RNA degradation machinery could modulate BP–3′-end distance and that introns with long BP–3′SS distances could produce 3′-trimmed isoforms that accumulate as stable introns. All tested 5′SS mutations decreased stable-intron accumulation (Fig. 5d and Extended Data Fig. 8c,d). The undetectable accumulation of the variant containing the GUAgua 5′SS from the *MER3* intron, which was the furthest from the consensus 5′SS sequence, could be attributed to the extremely inefficient splicing of this variant, as indicated by the greatly diminished lariat intron accumulation in the *dbr1Δ* mutant (Extended Data Fig. 8c). However, the decrease in accumulation of the other mutant introns could not be solely explained by decreased splicing. The GUAcGU 5′SS is the most common non-consensus 5′SS sequence, and splicing of this reporter variant was not significantly different from that of the wildtype variant (Extended Data Fig. 8c), consistent with the finding that introns containing this sequence do not exhibit slower splicing rates in vivo^56^. Nonetheless, this variant intron accumulated to a lower level than the wildtype intron, indicating that stable-intron accumulation is more sensitive to 5′SS mutation than is pre-mRNA splicing. Mutations to the BS also resulted in a significant, albeit more modest, reduction of stable-intron accumulation (Fig. 5e, Extended Data Fig. 8e,f). Together with the U1 snRNP and E-complex mutant phenotypes, our results showed that the U1 snRNP and 5′SS, two major determinants of spliceosome assembly, strongly contributed to the accumulation of stable introns, and perhaps to a lesser extent, the BS also contributes. These findings support the spliceosome reassembly model of stable intron accumulation, and not the reverse-splicing model.

### The spliceosome reassembles on linear excised introns in cells

The dependence of stable intron accumulation on E complex assembly factors and on strong 5′SS and BS sequences strongly supports the reassembly model for formation of the stable-intron complex. However, the ultimate test of the reassembly model is to examine whether a spliceosome can assemble and form a stable-intron complex on a linear RNA that has the same sequence as an authentic stable intron but was not generated by the spliceosome. We performed this test by using T7 polymerase to express a linear RNA with the same sequence as the MS2-tagged SAC6 stable intron in yeast cells. This artificial intron, expressed from an integrated, T7-driven cassette, contained a 5′ end defined by the T7 transcription start site and a 3′ end defined by the hepatitis delta virus (HDV) self-cleaving ribozyme (Fig. 6a) and thus was expected to differ from an excised intron at its ends (possessing a triphosphate rather than a monophosphate at its 5′ end and a cyclic phosphate rather than 2’, 3′ hydroxyls at its 3′ end). Nonetheless, this artificial intron accumulated as a full-length RNA species, and it did so more in saturated cultures than in log-phase cultures, mirroring the accumulation pattern of spliceosome-derived stable introns (Fig. 6b).

**Fig. 6:**
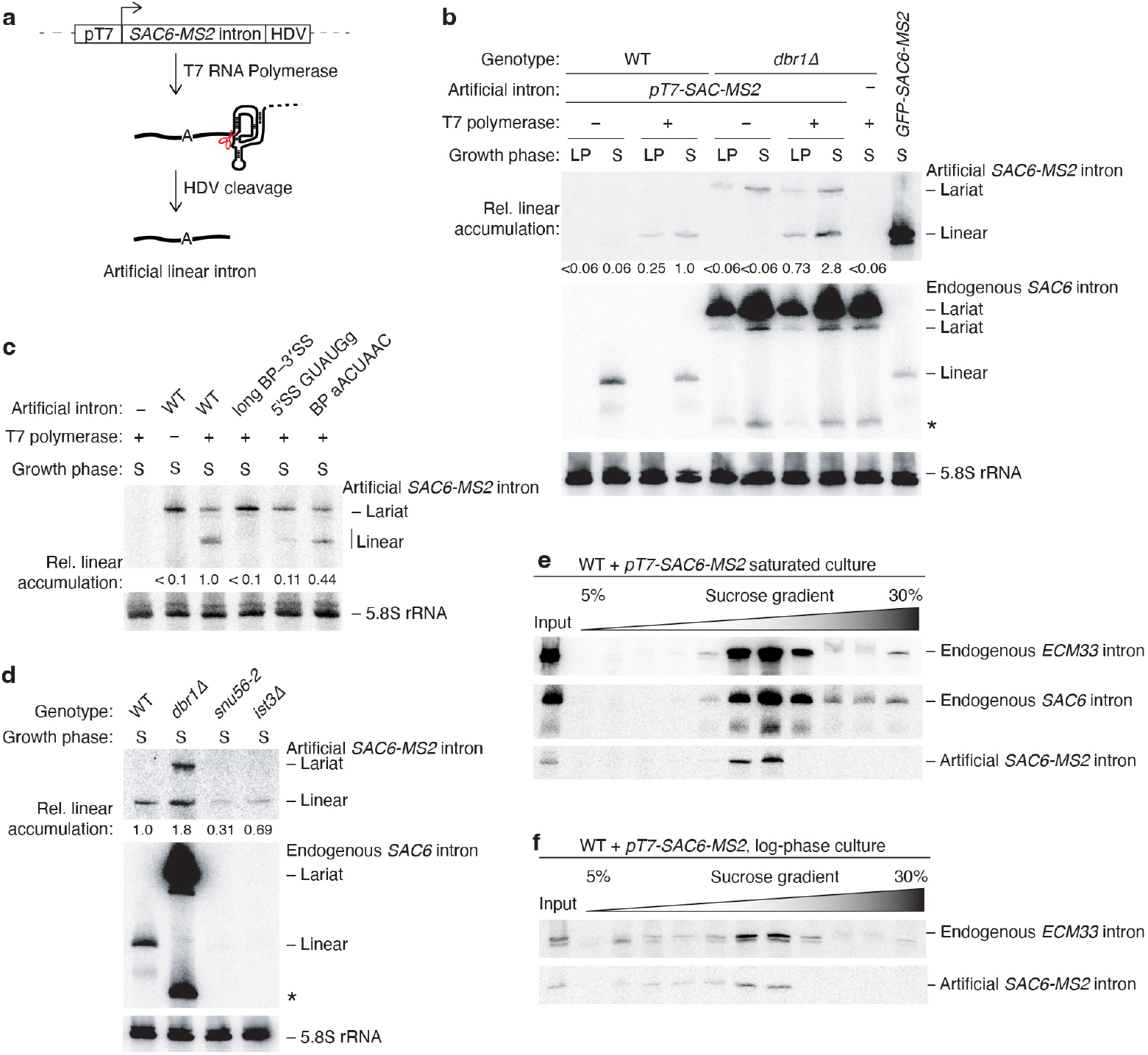
The spliceosome assembles on artificial linear introns. **a**, Design of artificial intron expression construct. The *SAC6-MS2* intron sequence is transcribed by T7 polymerase. Transcription begins at the first G of the intronic 5′SS sequence. HDV ribozyme self-cleavage defines a precise 3′ end. **b**, Accumulation of linear artificial introns in both wildtype and *dbr1Δ* strains. Shown is an RNA blot of a denaturing gel resolving total RNA from wildtype (WT) and *dbr1Δ* strains that contain the artificial intron construct (Fig. 6a), grown to either log-phase (LP) or saturation (S). Each strain additionally expressed either T7 RNA polymerase (+) or an empty-vector control (–). Blots were probed for the artificial intron using a probe that hybridizes to the MS2 tag (top) and then were reprobed for the endogenous *SAC6* intron (middle) and 5.8S rRNA (bottom; a loading control). The *SAC6* intron probe hybridized to both the artificial intron and the endogenous *SAC6* intron. For clarity, only the bands matching the migration of the endogenous intron are displayed. The migration of the linear artificial intron matched that of the *SAC6*-MS2 stable-intron spliced from *GFP* exons (rightmost lane). The slower-migrating lariat intron that accumulated in the *dbr1Δ* strain was dependent on integration of the artificial expression construct and independent of T7 polymerase expression. A band suspected to be a lariat degradation intermediate is also indicated (*). Numbers below the MS2 blot indicate the mean artificial intron level normalized to the WT strain with T7 polymerase after first normalizing all lanes to the 5.8S rRNA loading control. Blot is representative of *n* = 5 biological replicates. **c**, Dependence of artificial-intron accumulation on sequence features known to affect stable-intron accumulation. Shown is an RNA blot of a denaturing gel resolving total RNA from saturated-culture (S) *dbr1Δ* strains expressing T7 RNA polymerase and a series of artificial intron variants. The blot is otherwise as in **b**. *n* = 3 biological replicates. **d**, Dependence of artificial-intron accumulation on components of U1-snRNP and the RES complex. Shown is an RNA blot measuring the artificial intron accumulation in mutants that do not accumulate stable introns. Otherwise, this blot is as in **b**. *n* = 3 biological replicates. **e**, Co-sedimentation of the artificial intron with endogenous stable introns. Cleared lysate from a saturated culture of the wildtype strain transformed with the artificial intron and with the T7 polymerase plasmid was sedimented through a 5–30% sucrose gradient. Shown is an RNA blot of a denaturing gel that resolved an input sample representing 5% of the total lysate RNA and 25% of the RNA from each fraction oriented from low to high sucrose. The blot was probed for the *ECM33* intron (top), then stripped and reprobed using a probe that recognized both the endogenous *SAC6* intron (middle) and artificial MS2-tagged intron (bottom). **f**, Cosedimentation of excised introns and an artificial intron with high-molecular-weight complexes from log-phase cultures. Lysate from a log-phase culture of an artificial-intron-expressing wildtype strain was sedimented through a 5–30% sucrose gradient; otherwise, as in **e**.

**Fig. 7:**
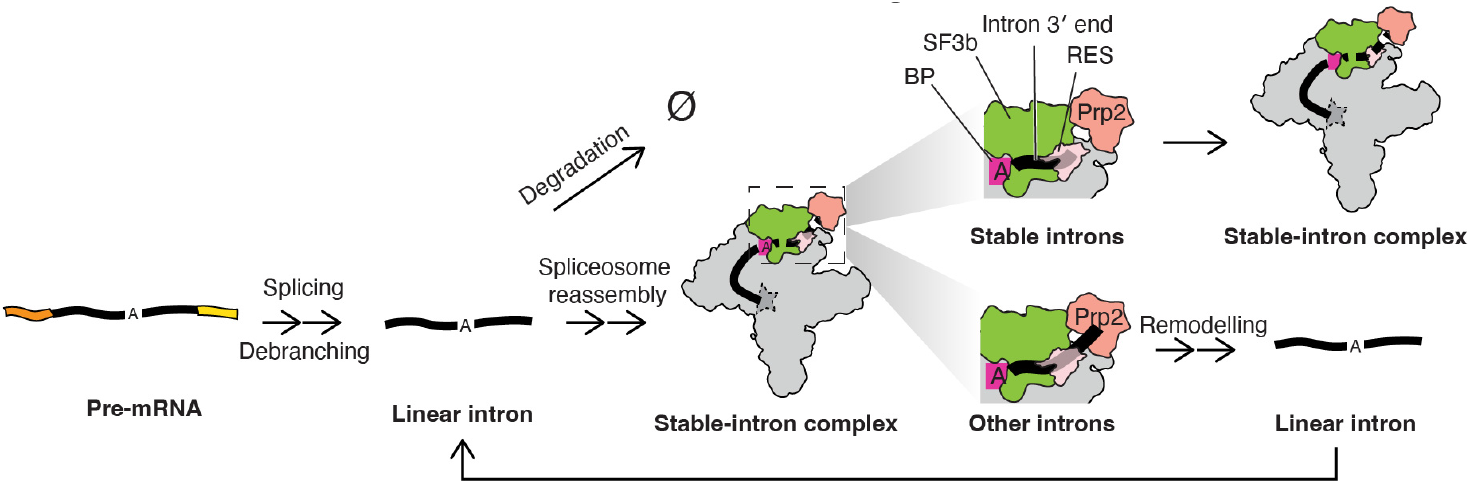
Model for the stabilization of excised linear introns. See main text for description

Because *DBR1* is required for the linearization of all introns excised by the spliceosome, including stable introns^1,55^, loss of the linear form and appearance of the lariat form in a *dbr1Δ* background provides a sensitive test for whether an intron was generated through the action of the spliceosome. When we examined artificial intron accumulation in a *dbr1Δ* background, we found that the abundance of the T7-polymerase-dependent linear species that accumulated specifically in saturated cultures did not decrease, which confirmed that this species was not a product of the spliceosome (Fig. 6b). Indeed, instead of decreasing, it increased > 2-fold. One way to explain this increase is that the artificial intron presumably competes with endogenous stable introns for access to the splicing machinery and that relief from this competition owing to the lack of endogenous stable introns supported greater accumulation of the artificial intron (Fig. 6b). A T7-polymerase-independent slower-migrating species was also observed. This species presumably corresponded to a lariat intron spliced from the product of non-specific Polymerase II transcription initiation upstream of the artificialintron-cassette integration site, as it was absent from a *dbr1Δ* strain that expressed T7 polymerase but lacked the artificial intron cassette (Fig. 6b). Although the lariat form of this spurious-transcription product was detectable in *dbr1Δ* cells, its linear form was not detectable in WT cells (Fig. 6b) and thus did not contribute to the signal observed for the T7-polymerase-dependent linear species, the accumulation of which strongly supported the reassociation model.

Further supporting the reassociation model, the T7-transcribed artificial intron and its derivatives recapitulated many key features of endogenous stable introns. First, as mentioned earlier, it accumulated more in saturated cultures than in log-phase cultures. Second, artificial intron accumulation depended on the same sequence features as endogenous stable introns; lengthening the BP–3′SS distance or mutating the consensus 5′SS and BS sequences reduced artificial intron accumulation (Fig, 6c and Extended Data Fig. 9a,b). Third, artificial-intron accumulation decreased in the U1-snRNP-mutant background *snu56-2* and RES-complex mutant background *ist3Δ* (Fig. 6d and Extended Data Fig. 9c,d). Finally, in both wildtype and *dbr1Δ* backgrounds, the *SAC6* artificial introns co-sedimented with endogenous stable introns produced in the wildtype background, indicating that they were bound by a large complex with similar physical properties as the stable-intron complex (Fig. 6d and Extended Data Fig. 9h).

Further inspection revealed a cryptic 5′SS within the *SAC6* stable intron. However, experiments mutating the cryptic site showed this it did not confound our analyses (Extended Data Fig. 9a,b). Moreover, artificial versions of the *UBC4* intron and an *ACT1* intron mutant containing a shortened BP–3′SS distance^1^—neither of which had a cryptic 5′SS—accumulated as full length RNA species in a T7-polymerase-dependent, saturated-culture-dependent, BP–3′SS-distance-dependent, and *DBR1*-independent manner (Extended Data Fig. 9e,f). Together, these results showed that expression of a linear intron sequence from an orthogonal T7 promoter was sufficient to recapitulate the behavior of stable introns produced by the spliceosome, thereby ruling out the spliceosome backtracking model and strongly supporting the spliceosome reassembly model.

In the course of these experiments, we noted that for some stable introns, accumulation was observed in logphase cultures, albeit at substantially lower levels than in stationary-phase cultures, implying that spliceosome assembly on excised linear introns occurs to some extent in logphase cultures. Examples included the *SAC6-MS2* artificial intron (Fig. 6b), some endogenous stable introns, such as *ECM33* (Fig. 3c), and the *SAC6-MS2* stable-intron spliced from *GFP* exons (Extended Data Figs. 1b and 9c). To investigate if the spliceosome was indeed able to reassemble on excised introns in log phase, we examined the sedimentation pattern of these introns. Both endogenous and artificial introns migrated to heavy-sedimenting sucrose-gradient fractions, indicating that reassembly on excised introns is a basal activity of the yeast spliceosome, in that it occurs, at least to some extent, in unsaturated as well as saturated cultures (Fig. 6f and Extended Data Fig. 9i).

## Discussion

The structural, genetic and molecular results presented here reveal a surprising activity of the yeast spliceosome. It reassembles on excised linear introns, which can protect these introns from degradation. Our results support the following pathway for reassembly (Fig. 6g): All introns are first spliced via the canonical splicing mechanism. The 3′ exon present in pre-mRNAs bound by canonical B^act^ spliceosomes provides sufficient RNA downstream of the BP to permit Prp2-mediated remodelling of all spliceosomes bound to pre-RNAs, regardless of the intron BP–3′SS distance. After exon ligation and mRNA release, the lariat product of splicing is debranched and some fraction of the resultant linear intron is not degraded but is instead recognized by U1 snRNP and Mud2–Msl5 to initiate an additional round of spliceosome assembly. Following an initial set of canonical remodelling steps, spliceosomes bound to excised introns arrive at a B^act^-like complex that contains a non-catalytic U6 ISL conformation and an improperly formed active site.

In the B^act^-like stable-intron complex, Prp2, which requires a minimum number of nucleotides downstream of the BP to remodel the B^act^ spliceosome, discriminates stable introns from other introns; the short BP–3′SS distances of stable introns cannot support efficient Prp2 translocation, leaving them trapped in the B^act^-like conformation, whereas the long BP–3′SS distances of the remaining introns permit Prp2-mediated remodelling even in the absence of a 3′ exon. Spliceosome remodelling by Prp2 exposes these introns to release and/or decay, thus preventing their accumulation.

Our U6 ISL structure resembles that of the U6 ISL in structures of the human spliceosome stalled between B and B^act^ stages (pre-B^act-2^)^23,57^. This intermediate conformation is proposed to bridge the U6-snRNP-like structure achieved by the U6 ISL following U4 removal by Brr2 (pre-B^act-1^) and the catalytic U6 ISL conformation achieved in B^act^ (Fig. 2f)^24,57^. This resemblance suggests that active-site formation might be impaired during the stable-intron B-to-B^act^-like transition due the absence of a covalently linked 5′ exon.

In addition to being recalcitrant to Prp2-mediated remodeling, the stable-intron complex must also resist the activities other DExD/H-box ATPases that help discard stalled spliceosomes assembled on defective substrates^10^. The B^act^-like conformation of the stable-intron complex may render it resistant to detection by proofreading DExD/H-box ATPases. For example, Prp16 and Prp22, which proofread the first and second catalytic steps of splicing, respectively, act on the 3′ exon or intron 3′ region and thus would also not be able to remodel stable-intron complexes that lack an accessible intron 3′ end^58,59^. Moreover, both helicases act by promoting spliceosome disassembly by Prp43, which appears to be to be unable to act before the action of Prp2^60^, consistent with structural data suggesting that Hsh155/SF3b1 and Prp45 in the B^act^ complex (and in the stable-intron complex) sterically occlude the binding sites for Ntr2, which is essential for recruiting Prp43 and its activating G-patch protein Ntr1 to the spliceosome^15,61–63^. How or if the stable-complex is disassembled upon re-entry into logphase remains to be investigated.

A surprising implication of the reassembly pathway is the notion that most introns remain susceptible to spliceosome binding after excision from their host pre-mRNA and thereby compete with bona-fide pre-mRNA substrates for spliceosome binding. In this pathway, even non-stable introns undergo futile cycles of binding and release by the spliceosome, which may further limit the effective cellular spliceosome pool during times of stress.

How might accumulation of spliceosome-protected introns be favored in some cell states but not others? Our detection of heavy-sedimenting excised introns in both logphase and saturated cultures suggested that the spliceosome is always able to protect some excised introns from degradation through the reassembly pathway. Perhaps excised introns are degraded more slowly in saturated cells, which could increase the frequency of spliceosome reassembly in these cells. Another nonexclusive possibility is that increased spliceosome availability in saturated cultures causes increased reassembly on excised introns. Indeed, the U1 snRNP increases in abundance during starvation conditions^64^, which combined with the decrease in the expression of intron-containing ribosome protein genes during saturated culture and other stresses^65^, suggest an increase in spliceosome availability, as has been suggested to explain the splicing of proto-introns with poor splice-site sequences during rapamycin treatment and meiotic mRNAs in sporulation conditions^66–68^. Sequestration of spliceosome factors by stable introns might serve to counteract an overabundance of spliceosomes, thereby protecting the transcriptome from the detrimental effects of mis-splicing during times of stress.

Although eukaryotic lineages outside of budding yeast typically possess an expanded set of spliceosomal proteins and splicing-related factors that aid recognition of degenerate splice-site sequences^8^, the requirements for spliceosome reassembly, namely introns that contain strong spliceosome-recruiting signals and spliceosomes that are competent to assemble on imperfect substrates, should be present in diverse eukaryotes. We therefore speculate that *S. cerevisiae* might not be the only species that has coopted spliceosome reassembly to regulate the splicing pathway.

## Acknowledgements

We thank members of the Bartel lab, G. Fink, J. Staley, and K.L.K. Pang for helpful discussions; D.H. Lin, A. Latifkar, and K. Xiang for experimental assistance; J.T. Morgan, R. Muller, J. Vilardell, K. Collins, T. Lu, R. Singer, I. Borodina and G. Fink for strains and plasmids; the Whitehead Genome Technology Core for sequencing; the Taplin Mass Spectrometry Facility for help with protein identification; S. Sterling, J. Podgorski, E. Brignole, and C. Borsa for the smooth running of the MIT.nano cryo-EM facility; T. Nanchung, N. Bonheur and the Glass Wash Core for lab materials. M.E.W. is grateful to Feng Zhang for funding and support. G.W.L. was supported in part by a scholarship from the Natural Science and Engineering Research Council of Canada. M.E.W. was an HHMI fellow of the Helen Hay Whitney Foundation. D.P.B. is an investigator of the Howard Hughes Medical Institute. This paper was typeset with the bioRxiv word template by @Chrelli: www.github.com/chrelli/bioRxiv-word-template

## Author contributions

G.W.L., M.E.W. and D.P.B. conceived of the project. G.W.L. and M.E.W. performed and analyzed the experiments. D.P.B. and M.E.W. supervised the project. G.W.L. wrote the original manuscript. G.W.L., M.E.W. and D.P.B. reviewed and edited the manuscript.

## Methods

### Yeast strains, genetic manipulations and plasmids

*S. cerevisiae* strains used in this study are listed in Supplementary Table 1. All strains were in the BY4741 background. Transformations were performed using standard methods^69^. The *bud13Δ, ist3Δ, snu56-2* and *CUT314Δ* strains were generated using a CRISPR-Cas9 system adapted for use in *S. cerevisiae*, using the entry plasmid pV1382 (a gift from Gerald Fink, Addgene plasmid # 111436)^70^. Clones were confirmed by colony PCR and Sanger sequencing, and counter selected for the Cas9 plasmid by plating onto 5-FOA media before their use in experiments. Oligonucleotides (Azenta) used for guide RNAs and repair templates are listed in Supplementary Table 2. Yeast Knockout (YKO) Collection strains (Horizon Discovery) were validated using colony PCR and Sanger sequencing. Plasmids were generated by HiFi Assembly (NEB) and/or site-directed mutagenesis using standard methods. The plasmids used in this study are listed in Supplementary Table 3. Independent transformants, or individual colonies in the case of YKO Collection strains, were used as biological replicates.

### Recombinant protein purification

MBP-MS2 was purified essentially as described^71^. pMBP-MS2 (a gift from Josep Vilardell, Addgene plasmid # 65104) was expressed in Rosetta2(DE3) *E. coli* (Novagen). Cells were grown to OD_600_ 0.5 in LB media, induced with 1 mM IPTG (GoldBio) for 3 h at 37ºC, lysed by sonication in buffer AB1 (20 mM HEPES-KOH pH 7.9, 200 mM KCl, 1 mM EDTA) with 1X cOmplete EDTA-free protease inhibitor cocktail (Roche) and clarified by centrifugation at 40,000*g* for 30 min. Clarified lysate was bound to Amylose High Flow Resin (NEB), washed with AB1, AB2 (20 mM HEPES-KOH pH 7.9, 20 mM KCl, 1 mM EDTA), and eluted with ABE (AB2 with 10 mM maltose). To remove contaminating *E. coli* RNA, fractions containing MBP-MS2 were bound to a Resource Q anion-exchange column (Cytiva) and eluted using a 20 mM–1M KCl gradient in buffer AB2. Eluted protein was concentrated and dialyzed into MCP storage buffer (20 mM HEPES-KOH pH 7.9, 50 mM KCl, 10% (v/v) glycerol, 0.2 mM EDTA, 0.5 mM DTT), aliquoted, flash frozen, and stored at –80ºC.

ProtA-PP7CP was purified essentially as described^72^. pETZZ-tevPP7CP(C68A, C73A) (a gift from Kathleen Collins, Addgene plasmid # 27548) was expressed in Rosetta2(DE3) *E. coli*. Cells were grown to OD_600_ 0.5 in 2XYT media, induced with 1 mM IPTG for 3 h at 37ºC and lysed by sonication in lysis buffer (20 mM HEPES pH 7.9, 1 mM MgCl2, 200 mM KCl, 10% (v/v) glycerol, 20 mM imidazole) with 1X cOmplete EDTA-free protease inhibitor cocktail and clarified by centrifugation at 40,000*g* for 30 min. Clarified lysate was bound to Ni-NTA agarose beads (QIAGEN), and beads were washed with Lysis buffer with 0.1% (v/v) IGEPAL CA-630 and then eluted (lysis buffer with 250 mM imidazole and 0.1% (v/v) IGEPAL CA-630). Eluted protein was purified using anion exchange, concentrated, dialyzed and frozen as described above.

### Cryo-EM sample preparation and data collection

Stable-intron complex purification procedure was adapted from protocols describing purification of spliceosomes from in-vitro splicing reactions using MS2-tagged substrates^18,71^ to purify MS2-tagged introns expressed endogenously in saturated cultures. The BY4741 strain was transformed with pGWL1188 (pTEF1-PP7-GFP-SAC6-MS2), a pRSII416-based^73^ centromeric, Ura-selectable plasmid that contained the *TEF1* promoter, 2XPP7 hairpins (derived from pDZ207, a gift from Robert Singer, Addgene plasmid # 35191) inserted 5 nt upstream of the start codon, the yeGFP ORF (a gift of Timothy Lu, Addgene plasmid # 64389), the *SAC6-MS2* intron placed 5 nt downstream of the start codon, and the *CYC1* terminator. The *SAC6-MS2* intron consisted of the *SAC6* intron sequence with 2XMS2 hairpins inserted 49 nt from the 5′SS and 43 nt from the BP. 2XMS2 hairpin sequences were based on previous efforts to purify stable introns^1^. Accumulation of the linear excised *SAC6-MS2* intron was confirmed by RNA blot (Extended Data Fig. 1b). The sequence of the *SAC6-MS2* intron is listed in Supplementary Table 4.

A single colony was grown overnight in five mL SC-Ura media, with 2% dextrose, expanded into a 500 mL culture and grown overnight, then seeded at OD_600_ 0.1 in 30 L of media and grown in 20 2.8L baffled flasks at 30ºC for 24 h, to a final OD_600_ of ∼6. The culture was pelleted, washed once with sterile water and once with HE-75 buffer (20 mM HEPES-KOH pH 7.9, 75 mM KCl, 0.25 mM EDTA, 5% (v/v) glycerol, 0.5 mM DTT). The ∼180 g wet-weight pellet was resuspended in 90 mL HE-75 with 1X cOmplete EDTA-free protease inhibitor cocktail (Roche) and 1X HALT protease and phosphatase inhibitor cocktail (Thermo) and frozen dropwise in liquid nitrogen. Pellets were lysed in a SamplePrep Freezer/Mill (8 cycles, 2 min on, 2 min off, 10 Hz). Lysate powder was thawed at 4ºC in a water bath and then clarified by centrifuging first at 25,000*g* for 32 min and then centrifuging the supernatant at 140,000*g* for 85 min. The clarified lysate was flash frozen in liquid nitrogen and stored at –80ºC.

4 nmol of MBP-MS2 and 4 nmol ProtA-PP7CP (a ∼30-fold molar excess over the amount of the *SAC6-MS2* stable intron in saturated cultures estimated from a quantitative RNA blot) were added to thawed, clarified lysate and incubated at 4ºC for 30 min. IGEPAL CA-630 was added to a final concentration of 0.05% (v/v), and 28 mL aliquots of this mixture were layered atop four 8 mL 40% glycerol cushions (HE-75 with 40% (v/v) glycerol, 0.05% (v/v) IGEPAL CA-630) and centrifuged at 144,000*g* (29,000 RPM) in a SW32-Ti rotor (Beckman) for 6.5 h. This step functioned to enrich high-molecular-weight spliceosome complexes and deplete endogenous RNAses and ribosomes. The high-glycerol cushion fraction was extracted and bound to 1.28 mL IgG Sepharose 6 Fast Flow affinity resin (Cytiva) overnight at 4ºC to deplete exon-containing spliceosomes. The next morning, the flowthrough from these beads was bound to 2 mL Amylose High Flow Resin (NEB) for 4 h at 4ºC, washed with 40 column volumes of HE-75 with 0.05% Igepal CA-630, and eluted with 5×1 mL maltose elution buffer (HE-75 with 0.01% (v/v) IGEPAL CA-630, 12 mM maltose). The fractions were analyzed using SDS-PAGE followed by silver staining, and the second and third fractions, which contained the bulk of the purified spliceosomes, were concentrated using a 100 kDa molecular-weight cutoff Ultra Centrifugal Filter (Amicon) and dialyzed against cryo-EM buffer (20 mM HEPES-KOH pH 7.9, 75 mM KCl, 0.25 mM EDTA) for 3 h using a 20-kDa molecular-weight cutoff Slide-a-Lyzer Mini (Thermo).

Freshly purified spliceosomes were applied to a glow-discharged (12 s at 25 mA) Cu300 R2/2 holey carbon grid with a 2-nm layer of amorphous carbon (Quantifoil) mounted in the chamber of a Vitrobot Mark IV (Thermo Fisher Scientific) maintained at 12°C and 100% humidity. After 30 s, grids were blotted for 2 s with Ø55 grade 595 filter paper (Ted Pella) at blot force setting 0 and plunged into liquid ethane. Cryo-EM data were collected using the Thermo Scientific Titan Krios G3i cryo TEM at MIT.nano using a K3 direct detector (Gatan) operated in super-resolution mode with 2-fold binning, and an energy filter with slit width of 20 eV. Micrographs were collected automatically using EPU in AFIS mode, yielding 48,858 movies at 130,000× magnification with a real pixel size of 0.663 Å. The exposure time per micrograph was 0.68 s, fractionated into 41 frames, with a total flux of 25.70 e^−^/pix/s giving a total fluence per micrograph of 40 e^−^/Å^2^. The nominal defocus range was –2.5 to –0.8 µm.

### Cryo-EM data processing

All cryo-EM data were processed using CryoSPARC (v4.7.0)^74^. Movies were corrected for motion with Patch Motion Correction with dose-weighting. CTF parameters were estimated using Patch CTF Estimation. Micrographs were denoised with the “Micrograph Denoiser” tool, using 100 training mics, and particles were picked on the denoised micrographs using the Blob Picker with 220 – 350 Å particle diameter and a 0.7-diameter minimum separation distance. Particles (2,057,201 total) were extracted with a 720-pixel box that was binned to 120 pixels and 3.978 Å/pix. One round of Heterogeneous Refinement was performed using consensus volumes of B^act^, C, and P-complex spliceosomes as references (EMD-30637, EMD-12107, and EMD-47157, respectively)^14,40,75^. Two copies of the B^act^ reference were provided. Interestingly, the single high-quality class emerged derived from the C-complex reference, but had B^act^-like features, whereas the B^act^ references resulted in low-quality reconstructions. Particles from this high-quality class (929,365 total) were re-extracted with a 720-pixel box binned to 480 pixels (0.9945 Å/pix) and refined with non-uniform refinement^76^ to 2.91 Å resolution (auto-tightened mask, FSC = 0.143). The particles were then subject to three rounds of heterogeneous refinement, using this high-quality reconstruction as a reference and the three low-quality reconstructions from the initial heterogeneous refinement as decoys. At each round, 613,248 (66%), 585,220 (95%), and 570,246 (97%) of particles produced a high-quality reconstruction, respectively, and were passed forward to the next round. Duplicate particles were then removed (300 Å minimum separation distance, keeping the particle with the best picking statistics). The resulting 522,517 particles were refined with non-uniform refinement, with dynamic masking disabled and with minimization over per-particle scale, to 2.81 Å resolution (auto-tightened mask, FSC = 0.143). To improve the resolution, micrographs were split into nine AFIS groups based on their beam shift values and each group was subject to two iterations of global CTF refinement, fitting tilt, trefoil, and anisotropic magnification. Non-uniform refinement then yielded a 2.74 Å resolution reconstruction (auto-tightened mask, FSC = 0.143). Reference-based motion correction ^77^ was then performed to produce 568,241 polished particles (1948 were rejected during motion correction) with a 720-pixel box (0.663 Å/pix). For subsequent steps, particles down-sampled to a 480-pixel box (0.9945 Å/pix) were used for refinements, with the 720-pixel particles used for reconstruction and sharpening after refinements. Particles down-sampled to a 120-pixel box (3.978 Å/pix) were used for classification steps.

The polished particles were refined with non-uniform refinement without dynamic masking to produce a 2.65 Å resolution reconstruction (auto-tightened mask, FSC = 0.143). Then 3D Classification without alignment was performed, with five classes and a filter resolution of 6 Å, using a focus mask on the core of the spliceosome that excluded SF3b, such that the mobility of SF3b wouldn’t influence the classification. Two of the classes were high-quality and represented two states of the spliceosome, while the other three classes were lower quality and each resembled one of these two states. The two high-quality classes (126,784 particles for “State I”, 153,189 particles for “State II”) were processed separately but identically. First, non-uniform refinement was performed without dynamic masking, giving 2.81 and 2.61 Å resolution reconstructions for states I and II, respectively (auto-tightened mask, FSC = 0.143). Next, per-particle defocus was refined with the “Local CTF Refinement” job, improving the resolutions to 2.67 and 2.49 Å, respectively. Then, local refinement was performed with a focus mask on the core, excluding SF3b, searching a range of 10 degrees and 5 Å, with pose/shift gaussian priors and re-centering of rotations and shifts each iteration. This gave core reconstructions at 2.58 and 2.44 Å resolution for states I and II, respectively. The same local refinement procedure was performed with a focus mask on SF3b, giving 2.53 Å and 2.56 Å-resolution SF3b reconstructions for each state. However, since no difference was observed in SF3b between the states (except for relative position to the core), the state I and II particles were merged and SF3b focus-refinement was repeated, producing a 2.41 Å reconstruction.

### Cryo-EM model building

For model-building and refinement, maps were reconstructed with a 720-pixel box (“Homogeneous Reconstruction Only” job) to produce smoother 0.663 Å/pix maps, which were cropped in real-space to 420 pixels to save memory during model building and refinement. The core maps were modelled by initiating with the core of the P-complex spliceosome (PDB 9DTR)^75^, as this was the highest-resolution catalytic spliceosome structure, with subunits not found in B^act^ removed. This structure was refined into the state I and state II core maps using ISOLDE^78^ with torsion and distance restraints, with the maps blurred in ChimeraX (v1.7–1.10)^79^ with a Gaussian filter and B-factor of 400 Å^2^ such that lower-resolution regions would be modelled accurately. The structures were then refined with Servalcat^80^ into cropped half-maps. Then, all side chains were inspected and fixed using Coot (v0.9.8)^81^ and peptides of Cus1, Bud13 (Cwc26), Hsh155, Prp11, and Cwc24 were built manually. Model-building was aided in cases of weak density with EMD-30637 (canonical B^act^ structure)^14^, provided that the lowpass filtered densities were consistent.

To build the SF3b lobe, an initial model was generated using AlphaFold3^82^, providing the full sequences of Hsh155, Cus1, Rse1, Hsh49, Ysf3, Rds3, Prp21, Prp11, Prp9, and Cwc24 as an input. The model was clustered into predicted aligned error (PAE) domains using ChimeraX, and each PAE domain was aligned to PDB reference 7DCO (canonical B^act^ structure)^14^ before merging, docking into the cryo-EM density, and restrained refinement using ISOLDE. Peptides with poor density were trimmed, and every side chain was inspected and manually adjusted using Coot. As above, model-building was aided in cases of weak density with EMD-30637 (canonical B^act^ structure), provided that the low-pass filtered densities were consistent. The Cus1/Hsh49 subcomplex had poor density in our map and in EMD-30637, so the crystal structure of this subcomplex (PDB 5LSL)^83^ was docked into the density and restrained in subsequent refinements. The RES complex (Bud13, also known as Cwc26, Pml1, Ist3) was co-folded with Prp45 in AlphaFold3 and docked and refined into the density.

After manual refinement, all models were refined into half-maps using Servalcat. Then, Q-scores^84^ were calculated, applied as attributes to the models within ChimeraX, and used to delineate well-resolved parts of the model (Q-score > 0.7). The well-resolved parts were refined manually using ISOLDE into sharpened density, whereas the less-well-resolved parts were refined into blurred maps (applying a Gaussian filter with B-factor of 200 – 400) with reference model restraints either from B^act^ (PDB code 7DCO) or P complex (PDB code 9DTR). Parameter files generated by ISOLDE were then used for restrained real-space refinement using Phenix^85^. The peripheral components of the spliceosome, including the U5 Sm ring, Prp19, Snt309, Cef1 C-terminus, Clf1 C-terminus, Syf1, Isy1, and Ntc20 were copied directly from P-complex (PDB code 9DTR) without refinement. ISOLDE was used with strong distance and torsion restraints to model the bend of the helical arches of Clf1 and Syf1 (and movement of the associated Isy1 and Ntc20) in B^act^ compared to P-complex. B-factors were not refined for these peripheral proteins. All Figures were generated using ChimeraX. Cryo-EM data statistics and model statistics can be found in Extended Data Table 1.

### Primer extension

RNA from aliquots taken over the course of the stable-intron complex purification protocol was extracted using TRI reagent (Invitrogen) according to the manufacturer’s protocol. The excised intron standard was generated by in-vitro transcription (HiScribe T7 High Yield RNA Synthesis Kit; NEB) of a SAC6-MS2 sequence that had a 5′ hammerhead ribozyme and 3′ herpes delta virus (HDV) ribozyme appended using two rounds of PCR. The premRNA standard was generated using a run-off in-vitro transcription of a T7 template generated through PCR of pGWL1188. Oligonucleotides used for generation of the template DNAs are listed in Supplementary Table 2. 10 fmol of standard RNA or total RNA equivalent to 0.007% of the input and exon-depleted samples or 0.035% of the cryo-EM sample were reverse transcribed using Superscript III (Life Technologies) according to the manufacturer’s protocol without dCTP and using a radiolabelled DNA primer that hybridized to the 5′-end of the SAC6 intron (oGWL1214; Supplementary Table 2). cDNA products were resolved on a denaturing poly-acrylamide gel and visualized using a Typhoon Phosphorimager (Cytiva).

### sRNA-seq

Stable-intron complexes were prepared using a version of the protocol detailed above that lacked the PP7-mediated exon-depletion steps. The strain used for this preparation was BY4741 transformed with pGWL1098 (pTEF1-URA3-ECM33-MS2) or pGWL1155 (pTEF1-URA3-SAC6-MS2). Both plasmids were based on Ura-selectable centromeric pRSII416 plasmids with the *URA3* promoter replaced by the *TEF1* promoter and either the *3P-ECM33* intron (pGWL1098; sequence in Supplementary Table 4 and Ref. *7*) or the *SAC6-MS2* sequence described above (pGWL1155) inserted 49 nt downstream of the *URA3* start codon.

Total RNA from these preparations was extracted using TRI reagent (Invitrogen), decapped and dephosphorylated using RNA 5′ Pyrophosphohydrolase (NEB), 5′ phosphorylated using PNK (NEB), and then used as input into the NEBNext Small RNA Library Prep kit (NEB) following manufacturer’s protocols. Libraries were sequenced on a MiSeq Nano with 125x125 bp paired-end reads. Adapters were trimmed using cutadapt^86^ (v4.8) with the parameters “--trim-n -q 20 -m 10 -a AGATCGGAAGAG-CACACGTCTGAACTCCAGTCAC -A GATCGTCGGACTG-TAGAACTCTGAACGTGTAGATCTCGGTGGTCGCCGTATCATT”, and reads were mapped using STAR^87^ (v2.7.1a) by first mapping to the MS2-intron plasmid sequence, and then mapping the remaining reads to the *S. cerevisiae* genome (R64-1-1, downloaded from www.yeastgenome.org)^88^ using the parameters “--alignIntronMax 1000 --sjdbOverhang 100 -- outSAMtype BAM SortedByCoordinate --quantMode GeneCounts --outReadsUnmapped Fastx” and with “–sjdbGTFfile” supplied with transcript annotations. Intron annotations were constructed with the *S. cerevisiae* genome features annotations (www.yeastgenome.org) and the Ares Lab Yeast Intron Database^89^ (http://intron.ucsc.edu/yeast4.3/). Reads were assigned using featureCounts (v2.0.1)^90^ to genes using annotations downloaded from the Saccharomyces Genome Database (vR64-1-1), introns using annotations constructed with the Ares lab database, and uncharacterized transcripts using annotations generated by Kang and colleagues that were modified to collapse overlapping features^91^. Counts for the endogenous and MS2-tagged *ECM33* and *SAC6* introns were summed. RNA-seq browser tracks were visualized using IGV (v2.18.2)^92^. Sequence logo was made with Logomaker^93^.

### Growth conditions and harvesting for RNA blots and RNA-seq

Yeast for RNA analysis were grown at 30ºC on standard synthetic complete (SC) dropout plates and media (Sunrise Biosciences) supplemented with 2% dextrose. SC-Ura and SC-His were used to maintain strains that contained plasmids that contained *URA3* or *HIS3-*marked plasmids. For each sample, a single colony was grown overnight in 5 mL of media, back diluted 1:25 to grow for another night (final OD_600_ ∼ 4–6), then used to seed a culture to OD_600_ 0.2. Log-phase cultures were harvested during early log phase (OD_600_ ∼0.5) 4–6 h after seeding, depending on the growth rate of the strain. Saturated cultures were harvested 24 h after seeding (final OD_600_ ∼ 4–6). Strains with slower growth rates often saturated at lower optical densities, and these strains were verified to have reached saturation by extending growth to 48 h and confirming that there was no additional increase in OD. All cultures were harvested by vacuum filtration and flash frozen in liquid nitrogen. Frozen pellets were mixed with frozen TES buffer (20 mM Tris pH 8, 1 mM EDTA, 1% (v/v) SDS; 400 µL per 50 OD harvested) and mechanically lysed using a Sample Prep 6870 Freezer/Mill Spex SamplePrep; 8 cycles of 2 min on, 2 min off at 10 Hz). Lysate powder was stored at –80ºC.

### RNA blots

Total RNA was extracted from frozen TES lysate powder or from aliquots of the stable-intron complex purification steps using TRI reagent (Invitrogen) according to the manufacturer’s protocol. P32-labelled Century markers (Ambion) and 15 µg of total RNA for each sample were resolved on a denaturing urea-polyacrylamide gel and transferred onto a BrightStar-Plus Positively Charged Nylon Membrane (Invitrogen) using a semi-dry transfer apparatus. For experiments that measured stable-intron accumulation, EDC (N-(3-dimethylaminopropyl)-N′-ethylcarbodiimide; Sigma-Aldrich) was used to chemically crosslink 5′ phosphates to the membrane^94^. For blots that included analyses of RNA species that did not possess 5′ phosphates (mRNAs, snRNAs, lariat introns and artificial introns), UV crosslinking was used instead (120 J/cm^2^). Blots were hybridized to radio-labelled DNA probes (Supplementary Table 2). A detailed protocol is available at http://bartellab.wi.mit.edu/protocols.html. RNA-blot data were analyzed with ImageQuant (v10.2). For experiments performed with multiple biological replicates, a single representative blot is shown, along with the mean quantified levels.

### RNA-seq

Total RNA was extracted from frozen lysate powder as described above. 0.1 ng of an equimolar mix of an in-vitro transcribed GFP, Renilla luciferase, and Firefly luciferase RNA mix was added to 1 µg of total RNA, which was then subjected to rRNA depletion using the riboPOOLs rRNA depletion kit (siTOOLs) according to the manufacturer’s protocol. RNA-seq libraries were prepared using an in-house protocol optimized for pairing to sequencing of ribosome-protected fragments, which reduced size bias^95^. rRNA-depleted RNA was fragmented, size-selected for 27–40-nt fragments, sequentially 3′ and 5′-ligated, reverse transcribed, and amplified using PCR. A detailed protocol is available at http://bartellab.wi.mit.edu/protocols.html.

### RNA-seq analyses

Reads were trimmed using cutadapt (v4.8) using parameters “-u 4 -m 10 -O 7 -a N{4}CTGTCTCTTATACACATCTCCGAGCCCACGAGAC” and mapped to the *S. cerevisiae* genome (R64-1-1) using STAR (version 2.7.1a) as described above. Aligned reads were assigned to genes and introns using STAR quantMode. Introns known to contain snoRNAs (Ares Lab Yeast Intron Database) were excluded from analyses. Branchpoint–3′SS distance annotations were calculated using the Ares Lab Yeast Intron Database and were defined as the distance from the annotated BP-A to the annotated 3′SS G inclusive. 5′SS and branch site sequences were from the Ares Lab Yeast Intron Database. Genes and introns that did not have at least 10 reads in at least half of the libraries used in each analysis were excluded. Filtered counts were input to DESeq2 (v1.38.3)^96^ without lfcshrink() for count normalization and fold-change calculation between samples. Soft-clipped reads containing non-templated 3′ oligo(A) nucleotides were extracted from mapped BAM files using extractSoftclipped (https://github.com/dpryan79/SE-MEI). Changes in relative expression of the spliced, intron-retained, and stable-intron isoforms were quantified using the Mixture-of-Isoforms (MISO, v0.5.4)^97^ framework.

Stable-intron identification was performed essentially as described previously^1^ using DESeq2-normalized count data from two biological replicates, each of log-phase and saturated cultures for each genotype analyzed. To be identified as a stable intron, the intron had to exceed the following thresholds. First, intron accumulation (normalized reads) in saturated culture had to be greater than half of its host mRNA accumulation. Second, intron accumulation in saturated culture had to be greater than twice that of its accumulation in log phase. Third, the change in the intron-to-mRNA ratio observed between saturated and log-phase cultures (intron accumulation index) had to be greater than four. Fourth, the intron must have ≥ 2 terminal adenylated reads mapping to the 3′ end of the intron across the two biological replicates of the saturated-culture condition, or MISO-based support for preferential accumulation of the stable-intron isoform in saturated culture as compared to log phase (Bayes factor ≥ 30 in at least one biological replicate).

### Intron variant reporter experiments

The wildtype *SAC6-MS2* intron reporter was constructed by replacing the pTEF1-PP7 portion of pGWL1188 with the *ACT1* promoter to form pGWL1274 (pACT1-GFP-SAC6-MS2). All other variants of the *SAC6-MS2* intron were constructed by site-directed mutagenesis of pGWL1274. To form the *UBC4-MS2* and *ACT1-MS2*-intron reporter plasmids, the *SAC6-MS2* sequence was replaced with *UBC4-MS2* (*UBC4* intron with 2XMS2 hairpins inserted 42 nt downstream of the 5′SS and 27 nt upstream of the BP) or *ACT1-MS2* (*ACT1* intron with 2XMS2 hairpins inserted 86 nt downstream of the 5′SS and 179 upstream of the BP) to form pGWL1354 (pACT1GFP-UBC4-MS2) and pGWL1356 (pACT1-GFP-ACT1-MS2), respectively. The BP–3′SS distance of the *UBC4-MS2* intron was increased from 26 nt to 45 nt to form UBC4-MS2^long^ (pGWL1355) and the BP–3′SS distance of the *ACT1-MS2* intron was decreased from 44 nt to 25 nt to form ACT1-MS2^short^ (pGWL1357). Intron sequences are listed in Supplementary Table 4. Variant plasmids were introduced into wildtype and mutant strains and selected for on SC-Ura plates. Individual colonies were grown, harvested, and analyzed by RNA blots as described above.

### Artificial-intron experiments

To form the artificial intron expression cassette (pGWL1322; pT7-SAC6-MS2-HDV), an EasyClone 2.0 genomic integration vector pCfB2337 (a gift from Irina Borodina, Addgene plasmid # 67555)^98^ that had been modified to remove any T7 promoter sequences was further modified to add the following components: 1) the T7 promoter sequence that lacked the terminal G bases, 2) the *SAC6*-MS2 intron sequence amplified from pGWL1274, 3) an HDV ribozyme sequence validated for used in yeast cells^99^, and 4) the T7 terminator sequence. This plasmid was the basis for all artificial intron variant plasmids. Plasmids were digested with NotI-HF (NEB), transformed into yeast and selected for using Hygromycin B (200 µg/mL) plate media. Integrants were confirmed by colony PCR. The artificial intron expression cassette sequence is listed in Supplementary Table 2.

To construct the T7 polymerase plasmid (pGWL1317; pTEF1-T7-Pol), the TEF1 promoter, a gBLOCK containing the T7 polymerase sequence (IDT), and the CYC1 terminator were inserted using Gibson Assembly into a His-selectable, centromeric pRS313 plasmid that had been modified to remove the T7 promoter sequence (pGWL1316; empty-vector control). These plasmids were transformed into the artificial-intron-cassette-containing strains described above, and transformants were selected by growth on SC-His media. Strains were analyzed by RNA blot as described above.

### Sedimentation velocity

Analysis of sedimentation patterns of stable and artificial introns was performed essentially as described^1^. 0.25 g flash-frozen yeast pellets were mixed with 1 mL frozen Lysis buffer (10 mM Tris-HCl pH 7.4, 5 mM MgCl_2_, 100 mM KCl, 1% Triton X–100, 1% Sodium Deoxycholate, 2 mM DTT, 20 U/ml SUPERasein (Life Technologies), 1X cOmplete EDTA-free protease inhibitor cocktail (Roche), 1X HALT protease and phosphatase inhibitor cocktail (Thermo), 100 ng/µL cycloheximide) and lysed in a SamplePrep Freezer/Mill (8 cycles, 1 minute on, 1 minute off, 10 Hz). Thawed lysate was clarified by centrifugation at 1,300*g* for 10 min, then 25 OD_260_ of clarified lysate was layered atop a 12.5 mL linear 5–30% sucrose gradient (20 mM HEPES-KOH pH 7.4, 5 mM MgCl_2_, 100 mM KCl, 2 mM DTT, 20 U/mL SUPERasein) and centrifuged in a SW-41 Ti rotor at 38,000 rpm for 4 h at 4ºC. Gradients were fractionated into 1 mL f
ractions using a Piston Gradient Fractionator (Biocomp). RNA from each fraction was extracted using TRI reagent, and RNA blots were performed as described above, except loading equivalent proportions of each gradient fraction instead of normalizing for RNA content. 4°C Gietz, R. D. & Schiestl, R. H. High-efficiency yeast transformation using the LiAc/SS carrier DNA/PEG method. *Nat. Protoc*. 2, 31–34 (2007).

**Extended Data Fig. 1:**
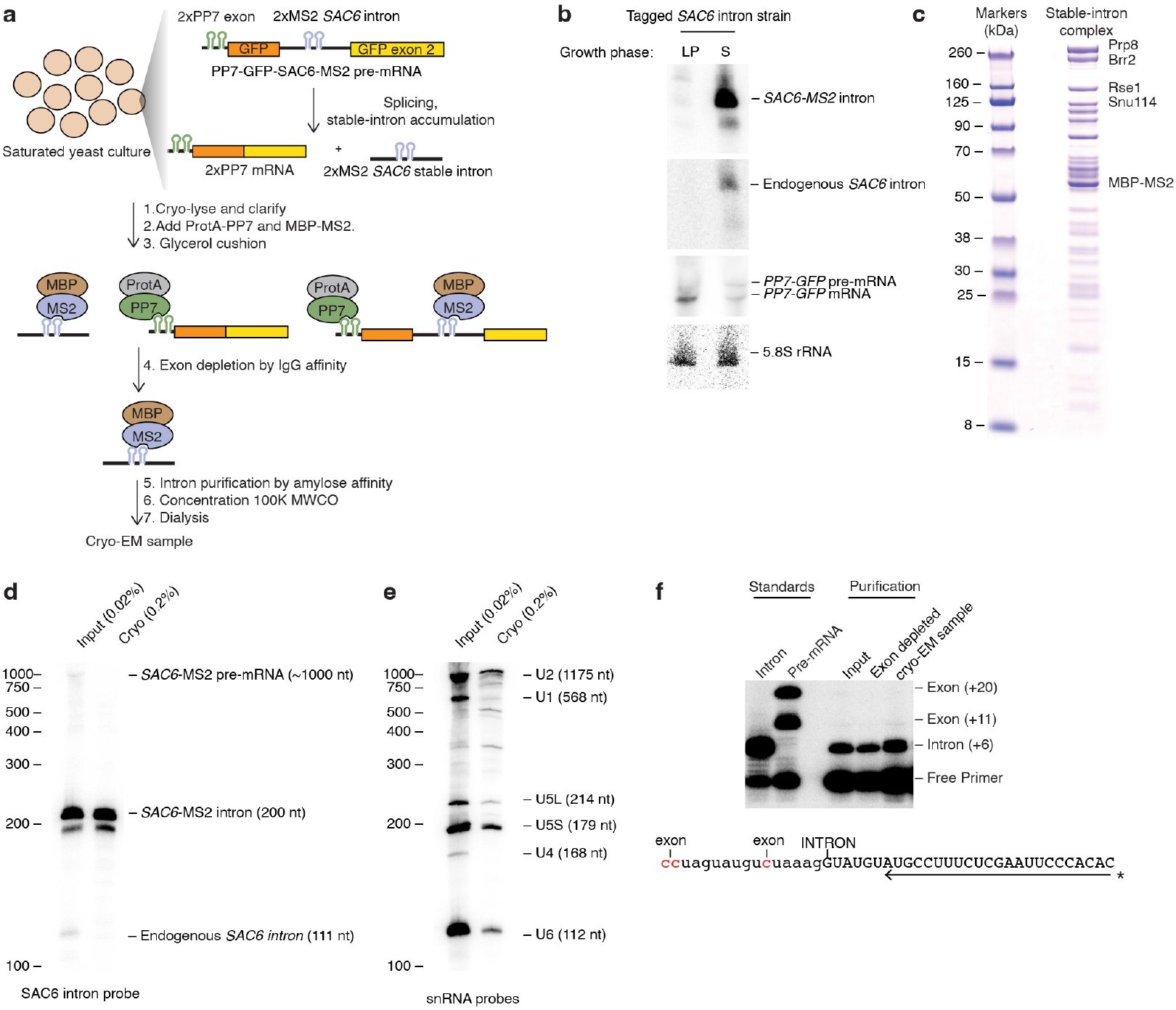
Purification of the stable-intron complex. **a**, Schematic for purification of the stable-intron complex from saturated cultures. MBP-MS2, Maltose-binding protein fused to MS2 coat protein; ProtA-PP7, 2X ProteinA fused to PP7 coat protein. **b**, Similar behaviors of tagged stable intron and endogenous stable intron. Shown is an RNA blot of a denaturing gel resolving total RNA from log-phase (LP) and saturated (S) cultures of a wildtype strain that expressed the *PP7-GFP-SAC6-MS2* pre-mRNA described in (A). The blot was probed for the MS2 tag (top), and then reprobed for the endogenous *SAC6* intron (upper middle), PP7 tag (lower middle), and 5.8S rRNA (bottom), which served as a loading control. **c**, Proteins in the stable-intron complex. Shown is an SDS-PAGE gel that resolves the cryo-EM sample, stained with Imperial Stain. Bands were labelled based on comparison to the distinct migration patterns of spliceosomal proteins observed in previous spliceosome preparations^18,100^. **d**,**e**, Enrichment of *SAC6-MS2* stable intron (**d**) and snRNAs (**e**) in purified stable-intron complex. Shown is an RNA blot of a denaturing gel resolving the indicated proportion of the glycerol cushion fraction (Input) and the concentrated and dialyzed sample used for cryo-EM analysis (Cryo). The blot was probed using an oligonucleotide that hybridized to a sequence common to the endogenous *SAC6* intron and the tagged *SAC6-MS2* intron (**d**), and then stripped and reprobed for U1, U2, U4, U5 and U6 snRNAs simultaneously (**e**). Migration of markers with lengths indicated (nucleotides) is at the left. **f**, No detectable upstream exon in the purified stable-intron complex. Shown is a Urea-PAGE gel resolving the products of primer extension across the *GFP-SAC6-MS2* exon–intron boundary at the 5′SS, performed in the absence of dGTP on RNA extracted from equal fractions of the glycerol cushion fraction (input) and IgG affinity flowthrough (exon depleted), and a five-fold fraction of the purified stable-intron complex (cryo-EM sample), as well as on in-vitro-transcribed RNAs representing the intron and the pre-mRNA (standards). Products terminating at the 5′-end of the excised intron (+6) represent the excised intron, and products terminating at the upstream C nucleotides of the 5′ exon (+11 and +20) represent the unspliced pre-mRNA. The +20-extension product was a result of reverse transcriptase readthrough of the first C at the +11 position, as verified by results from the in-vitro-transcribed standards. The in-vitro-transcribed intron standard consisted of the *SAC6-MS2* intron with a precise 5′-end defined by hammerhead ribozyme cleavage, and the pre-mRNA standard consisted of an RNA spanning 166 nucleotides upstream to 66 nucleotides downstream of the intron.

**Extended Data Fig. 2:**
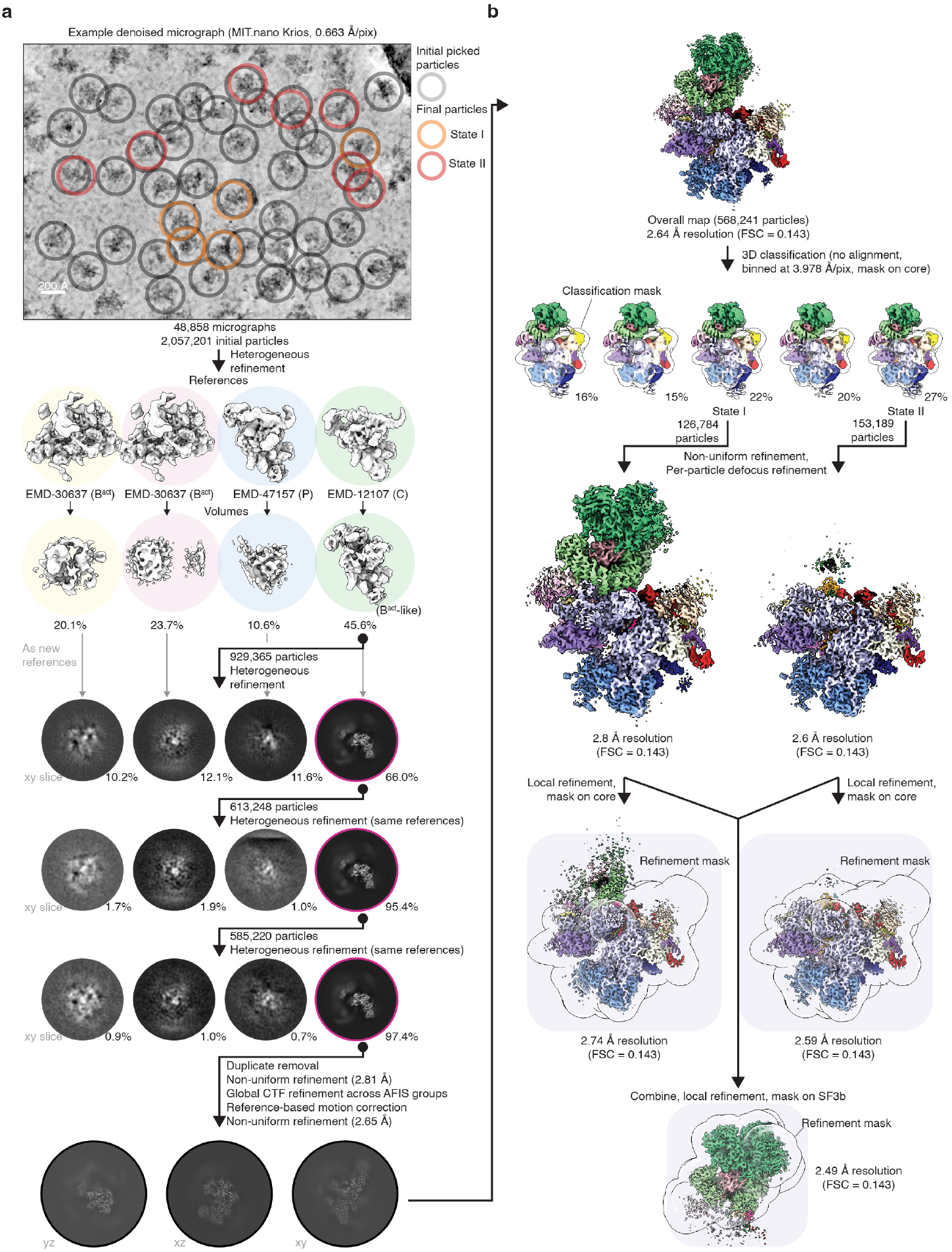
Collection and processing of the cryo-EM data. **a**, Representative micrograph (denoised using cryoSPARC) and heterogenous refinement scheme. Consensus volumes of B^act^ (two copies), C- and P-complex spliceosomes were provided as initial references. **b**, Focused classification and refinement scheme for the core and SF3b regions of the stable-intron complex.

**Extended Data Fig. 3:**
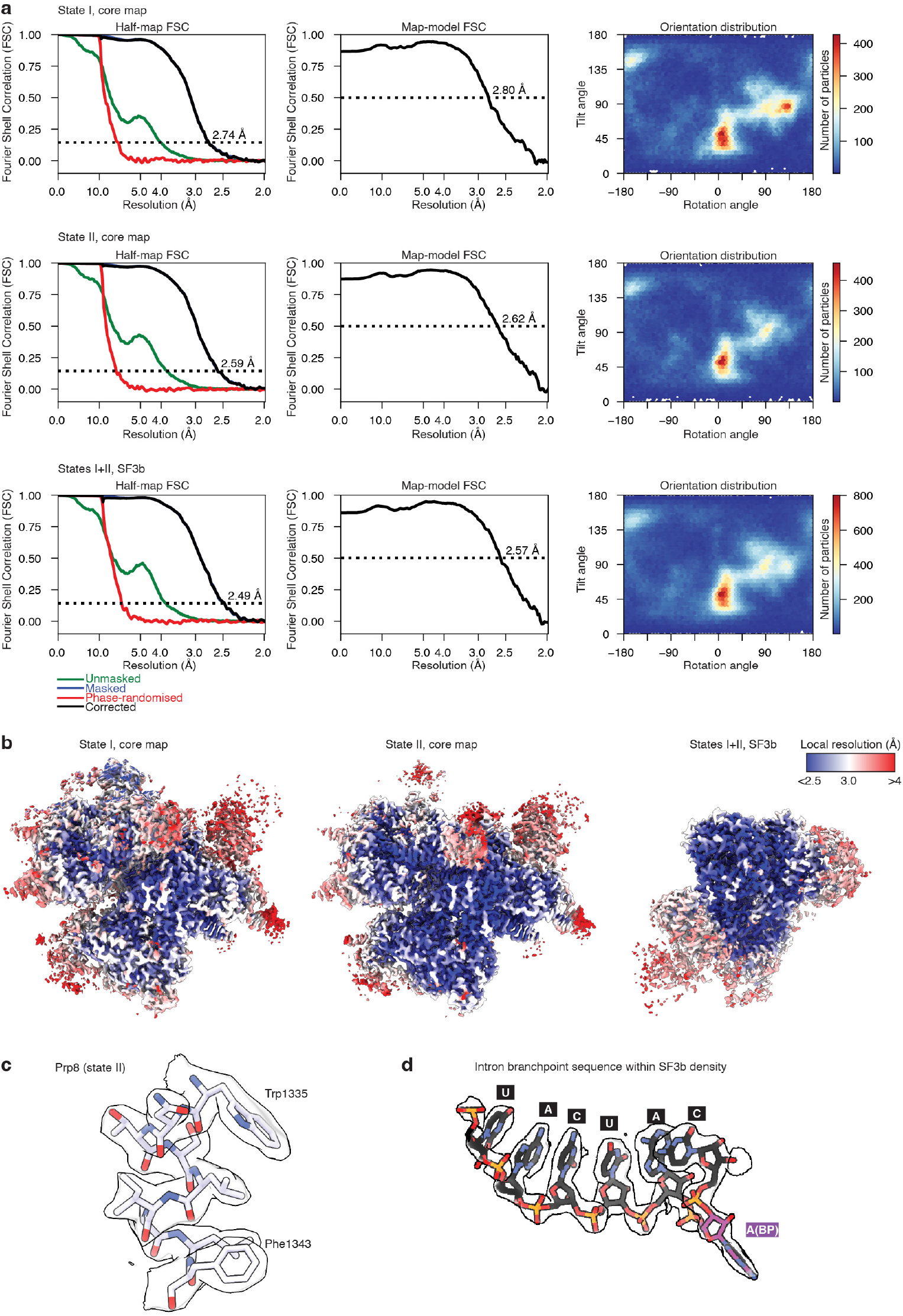
Cryo-EM density of the stable-intron complex. **a**, Gold-standard Fourier-shell correlation (FSC) curves (calculated with RELION), map-model FSC curves (calculated with PHENIX) and orientation distribution plots for the focus-refined maps. **b**, Focus-refined maps colored by local resolution (calculated using cryoSPARC). **c**, Density for Prp8 residues 1335–1343 in the state II core map. **d**, Density for the intron branch site in the SF3b map.

**Extended Data Fig. 4:**
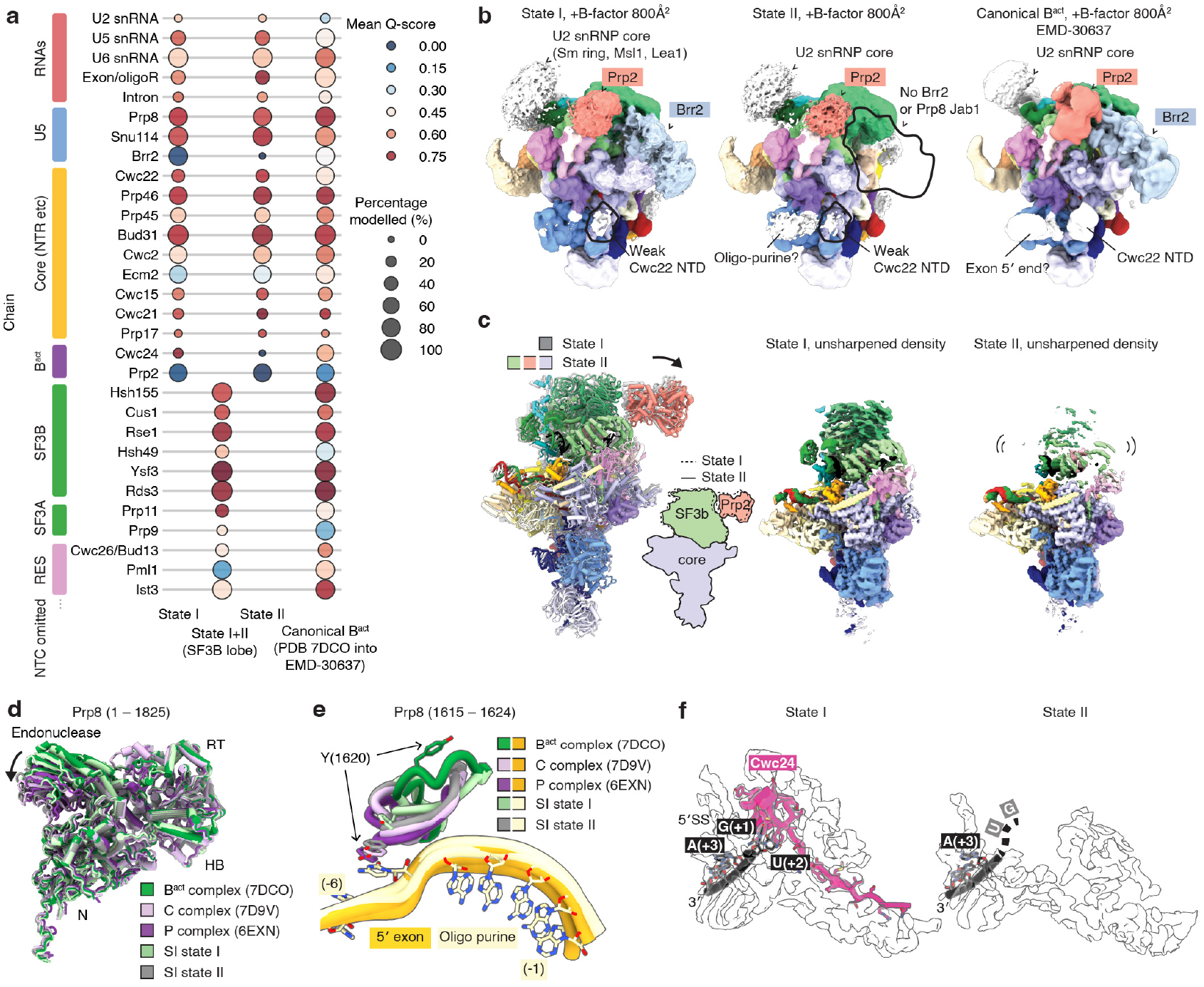
Differences between states I and II of the stable-intron complex and the B^act^ complex. **a**, Resolvability of modelled RNAs and proteins in the stable-intron-complex states I and II and the yeast B^act^ complex. Plotted are the mean Q-score, a measure of resolvability^84^, and fraction of modelled residues for each protein and RNA in the indicated complex. Because the SF3B, SF3A and RES-complex proteins were built using the focus-refined SF3B-lobe cryo-EM density shared by state I and state II (Extended Data Fig. 2), only one value is reported for each protein of this lobe. **b**, Low-pass filtered cryo-EM density maps of state I and state II of the stable-intron complex and the yeast B^act^ complex, colored by the associated models. Large-scale differences between the density maps are indicated using a black outline. **c**, Relative position and flexibility of the SF3B lobe of state I and state II of the stable-intron complex. Shown are the atomic models of the well-resolved core and SF3B lobe of the stable-intron complexes, and the unsharpened densities coloured according to the molecular models. The arrow indicates the shift in the position of the SF3b lobe in state II relative to state I. **d**, Shift of the Prp8 endonuclease domain towards a C- or P-complex-like position in state II of the stable-intron complex. Shown are molecular models of the Prp8 N-terminal domain (N), endonuclease domain, reverse transcriptase domain (RT) and helical bundle domain (HB) in the two states of the stable-intron complex and in the yeast B^act^, C and P complexes^14,40,75^. Models were aligned on Prp8. The arrow indicates the shift in the position of the Prp8 endonuclease domain in state II relative to state I. **e**, Shift in stable-intron complex Prp8 residues 1615–1624 towards the RNA positioned in the 5′-exon-binding site. For clarity, only the nucleobases from state II of the stable-intron complex are shown. **f**, Weak density for Cwc24 and the 5′ GU of the 5′SS in state II of the stable-intron complex.

**Extended Data Fig. 5:**
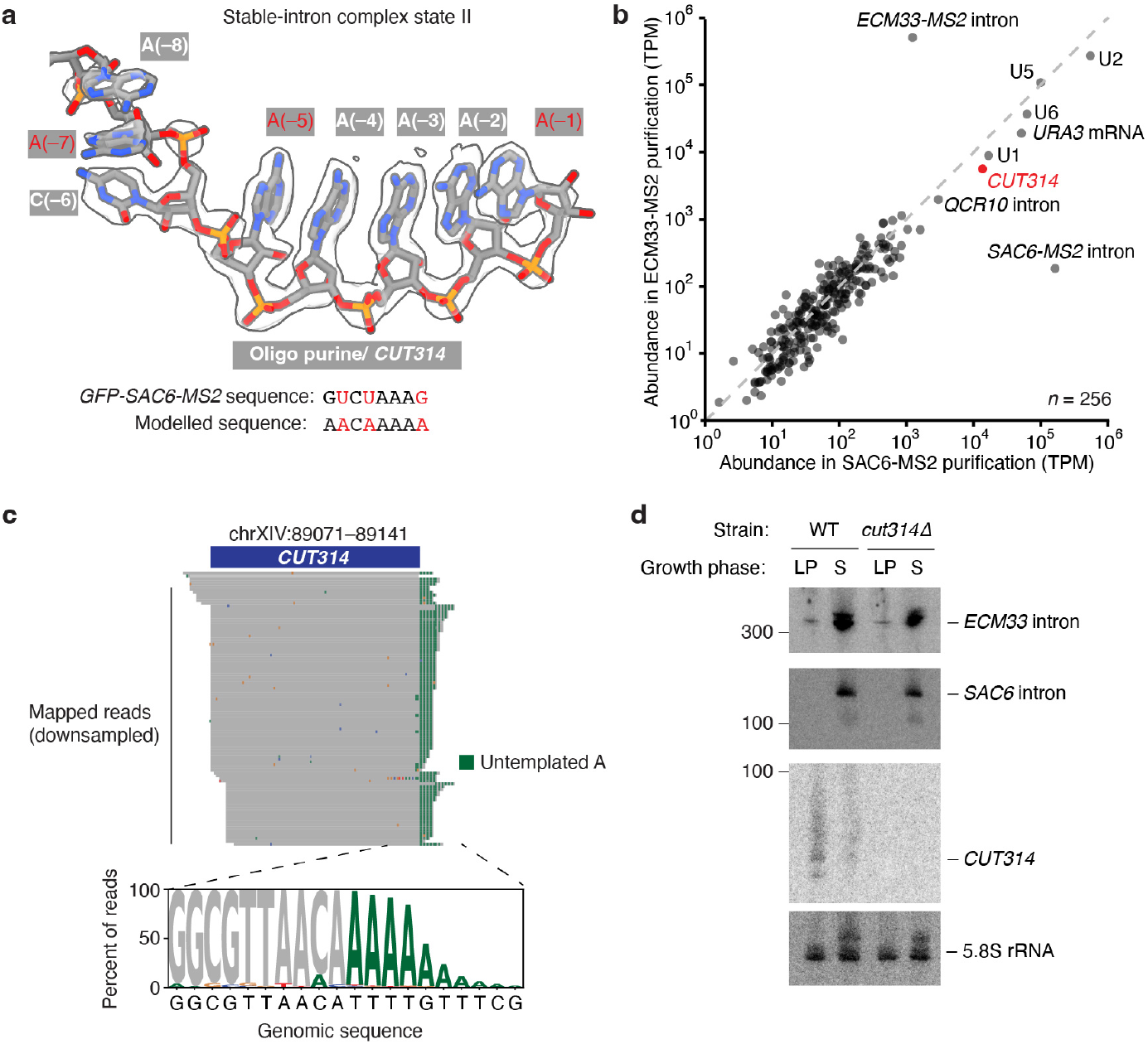
Copurification of CUT314 with the stable-intron complex. **a**, Sequence determination of RNA bound in the stable-intron complex using the stable-intron complex state II cryo-EM density map. Nucleotides at which the *GFP* 5′-exon sequence disagrees with the resolved density are labeled in red. **b**, RNAs observed in preparations of stable-intron complex. The scatter plot shows the abundance of RNAs observed in smaller-scale preparations of the *SAC6-MS2* and the *ECM33-MS2* stable-intron complexes, as determined by small RNA sequencing (sRNA-seq; expression cutoff, 1 transcript per million (TPM) in both samples). In these preparations, the stable introns were expressed from *URA3* exons, and an exon-depletion step was not implemented. *CUT314* RNA (red) was abundant in both preparations. **c**, sRNA-seq reads mapping to the *CUT314* locus. A representative downsampled set of 100 mapped reads are shown. The sequence logo was calculated from reads aligning to the 3′-end of the *CUT314* locus. **d**, No detectable effect of *CUT314* deletion on stable-intron accumulation. Shown is a blot of a denaturing gel resolving RNA from log-phase (LP) and saturated (S) cultures of the wildtype strain (WT) and a strain in which a genomic region spanning the full-length *CUT314* RNA was precisely deleted. The blot was probed simultaneously for the endogenous *ECM33* (top) and *SAC6* introns (second) and then stripped and reprobed for the *CUT314* RNA (third) and 5.8S rRNA (bottom), which served as a loading control. *n* = 1 biological replicate.

**Extended Data Fig. 6:**
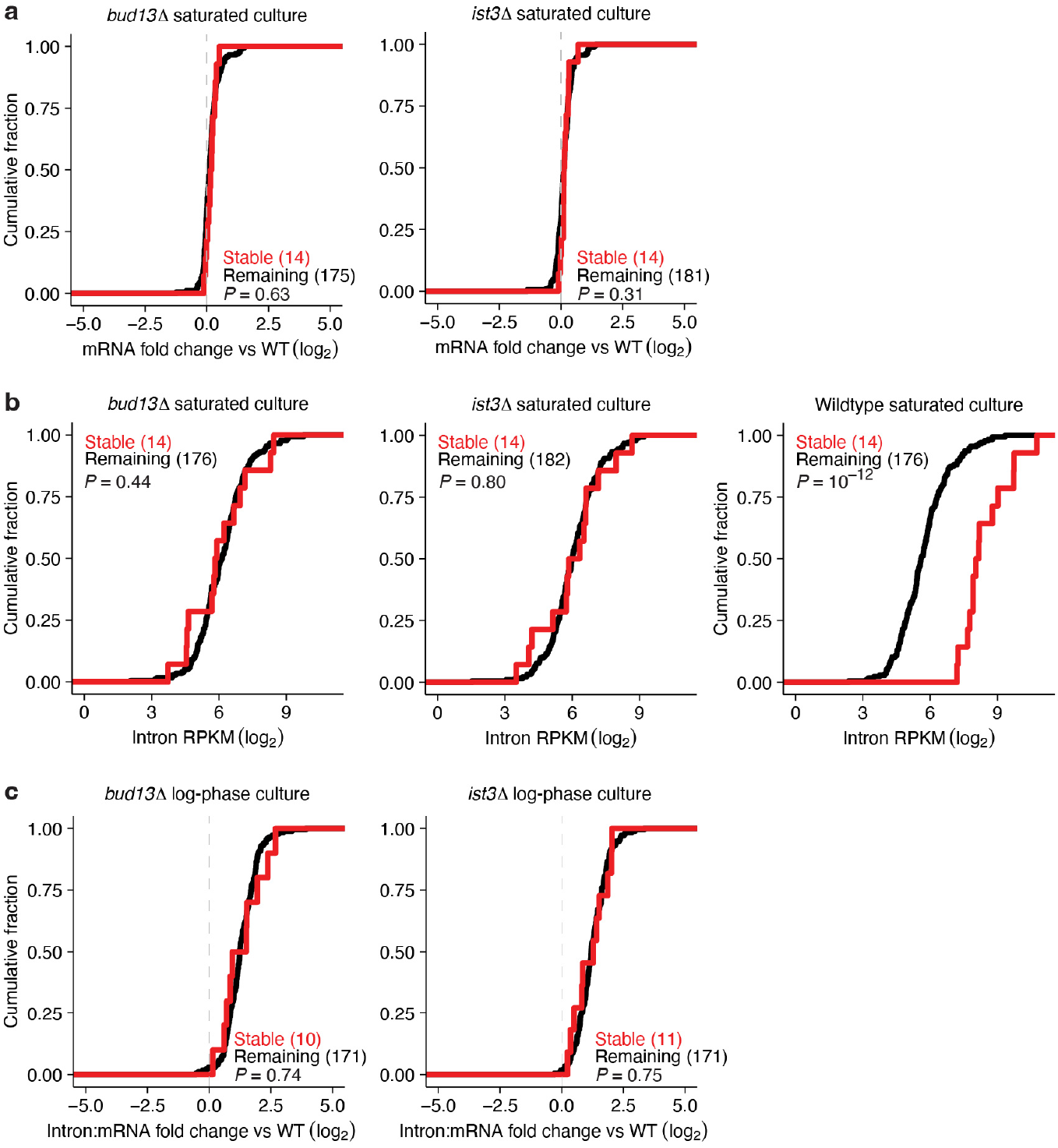
Mutants in the RES-complex cause stable introns to behave more like other introns. **a**, No detectable difference in mRNA levels observed between stable introns and other introns in RES-complex mutants in saturated cultures. Plotted are cumulative distributions of mRNA mean fold changes observed between saturated wildtype (WT) and saturated mutant cultures, comparing mRNAs that hosted stable introns identified in the matched WT sample (Extended Data Table 2) and those that hosted the remaining expressed introns. *P* values, two-tailed Kolmo-gorov–Smimov test. *n* = 2 biological replicates. **b**, No detectable difference in intron accumulation observed between stable introns and other introns in RES-complex mutants in saturated cultures. Plotted are cumulative distributions of mean length- and sequencing-depth-normalized intron read counts (RPKM), comparing results for stable introns defined in the matched WT sample (Extended Data Table 2) with results for other introns. At the left and center are results for RES-complex mutants, and at the right are results for matched WT cells. *P* values, two-tailed Kolmogorov–Smimov test. *n* = 2 biological replicates. **c**, No detectable difference observed between accumulation of stable-introns and other introns in RES-complex mutants during log phase. Plots are as in Fig. 3d,e, but for log-phase cultures. *P* values, two-tailed Kolmogorov–Smimov test. *n* = 2 biological replicates.

**Extended Data Fig. 7:**
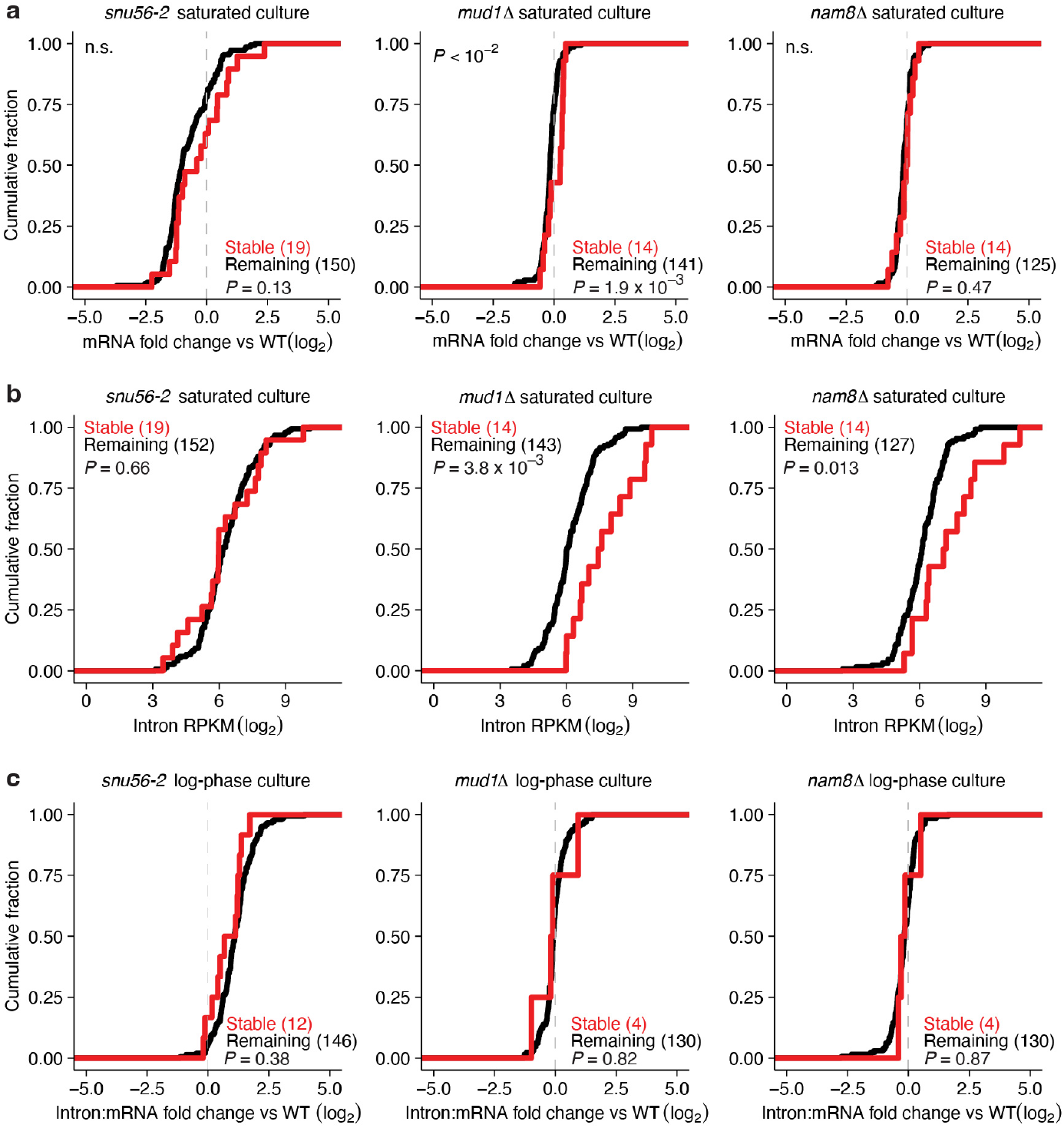
Mutants in U1-snRNP cause stable introns to behave more like other introns. **a**, No detectable difference in mRNA levels observed between stable introns and other introns in U1-snRNP mutants in saturated cultures; otherwise, as in Extended Data Fig. 6a. **b**, Smaller difference in intron accumulation observed between stable introns and other introns in U1 snRNP mutants, as compared to that observed in WT cultures (Extended Data Fig. 6c); otherwise, as in Extended Data Fig. 6b. **c**, No detectable difference observed between accumulation of stable introns and other introns in U1 snRNP mutants during log phase. Plots are as in Extended Data Fig. 6c.

**Extended Data Fig. 8:**
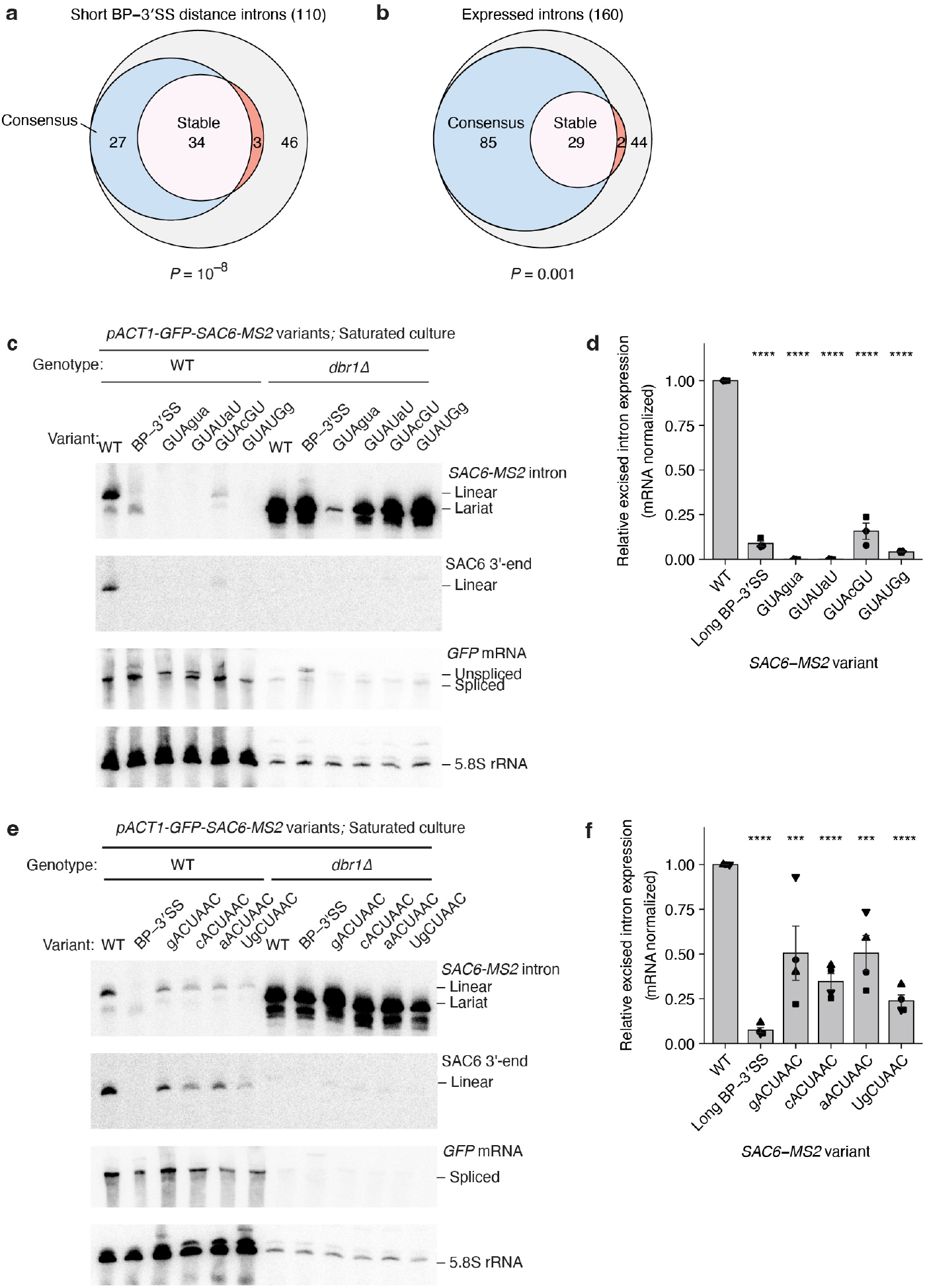
Effect of 5′SS and BS mutations on stable-intron accumulation. **a**,**b**, Enrichment of consensus 5′SS and BS sequences in stable introns compared to enrichment in other introns with short BP–3′SS distances (**a**) or other introns expressed in saturated cultures (**b**). Plots are as in Fig. 5a, except only analyzing either introns with BP–3′SS distances <30 nt—a cutoff that includes 37/41 stable introns (Extended Data Table 2) (**a**) or introns for which both the intron and the host mRNA met a 10-read cutoff in each of the wildtype saturated-culture sequencing libraries analyzed (**b**). *P* values, hypergeometric test. **c**,**e**, Reduced accumulation of introns containing 5′SS mutations (**c**) and branch site mutations (**e**). Shown are blots of denaturing gels resolving total RNA from saturated wildtype (WT) or *dbr1Δ* strains expressing the indicated *GFP-SAC6-MS2* variant. Blots were probed for the MS2 tag sequence (top), present in both the excised linear intron that accumulated in the WT strain and the lariat species that accumulated in the *dbr1Δ* strain. The blot was then sequentially stripped and re-probed for 1) the sequence near the 3′ terminus of the *SAC6* intron (which was shared between the WT sequence and the long BP–3′SS distance mutant) (upper middle), 2) the downstream exon of the intron-expression construct (designed to detect the *GFP* mRNA; lower middle), and 3) 5.8S rRNA (bottom). *n* = 3 (**c**), and *n* = 5 (**e**). **d**,**f**, Quantification of mRNA-normalized accumulation of introns with 5′SS mutations (**d**) or BS mutations (**f**). Plotted are the excised intron abundances of the variants tested in **c** and **e**, normalized to the WT intron sequence after first normalizing to the spliced *GFP* band. *n* = 3 (**d**), and *n* = 4 (**f**) biological replicates. Plots are otherwise as described in Fig. 5d.

**Extended Data Fig. 9:**
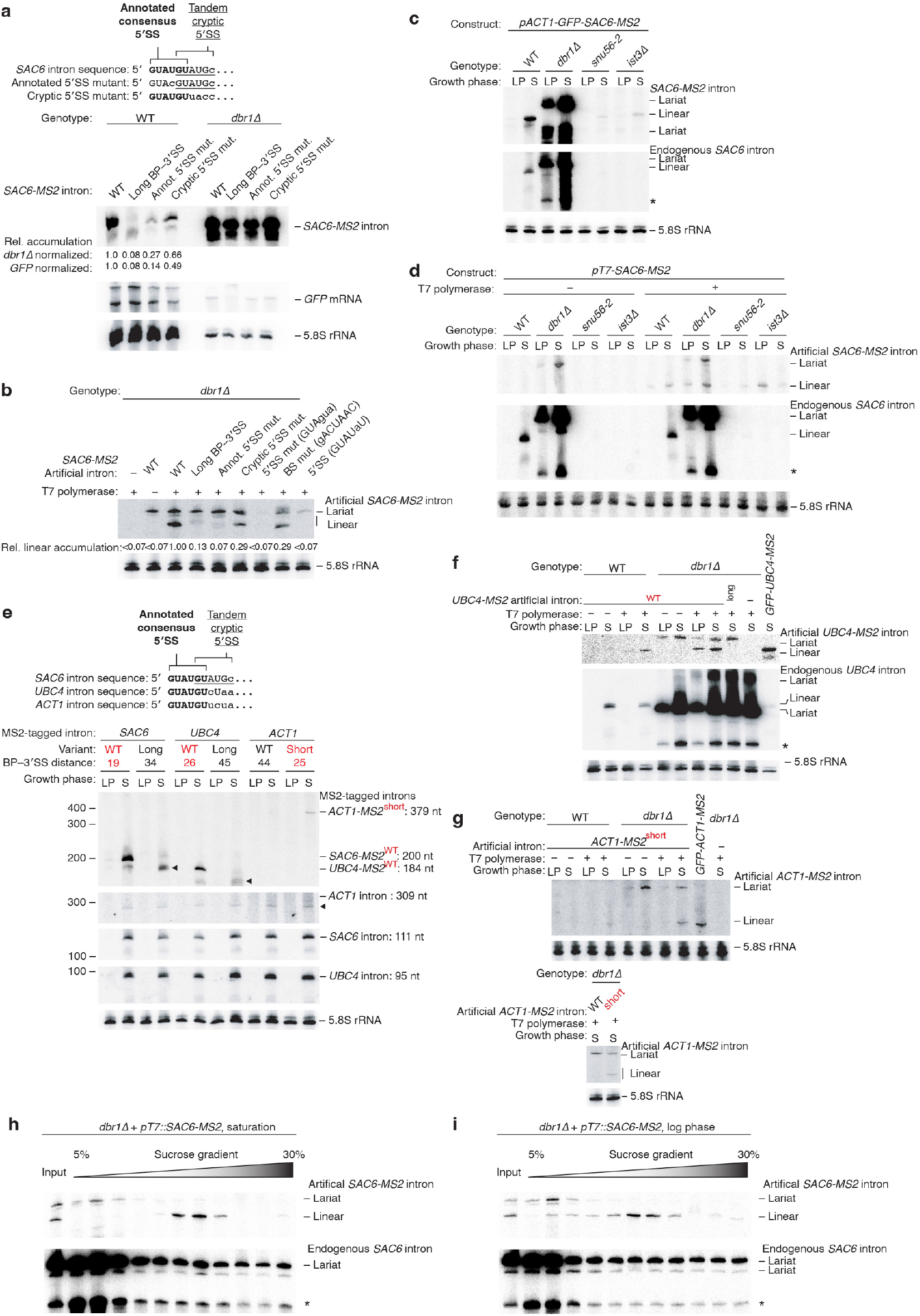
Effects of intron sequence and spliceosome mutation on artificial-intron accumulation. **a**, Greater contribution of annotated 5′SS than tandem-cryptic 5′SS to *SAC6* intron accumulation. The diagram (top) shows the 5′-terminal sequence of the *SAC6* intron, highlighting the tandem 5′SS and the mutations made to interrogate the relative contributions of the two 5′SSs. The representative RNA blot (below) is as described in Extended Data Fig. 8c. Also shown are the mean relative stable-intron accumulations from *n* = 2 biological replicates. **b**, Effect of mutations of the annotated 5′SS, the cryptic 5′SS and the BS on artificial intron accumulation. Annotated and cryptic 5′SS mutants are as described in **a**. Blot is otherwise as in Fig. 6c, showing mean relative accumulations from two biological replicates. **c**, Control for Fig. 6d showing dependence of *SAC6-MS2* stable intron accumulation on debranching enzyme and components of the U1-snRNP and the RES complex. Shown is an RNA blot resolving RNA extracted from log-phase (LP) and saturated (S) cultures of the indicated strains transformed with the *GFP-SAC6-MS2* reporter described in Fig. 5c. The blot was sequentially probed for MS2 tag (top), the endogenous *SAC6* intron (middle), and the 5.8S rRNA (bottom), which served as a loading control. A band suspected to be a lariat degradation intermediate is indicated (*). *n* = 3 biological replicates. **d**, Control for Fig. 6d showing dependence of artificial intron accumulation on T7-polymerase, saturated culture, and spliceosome components. Blots are as in Fig. 6b. *n* = 3 biological replicates. The reduced sensitivity to *SNU56* and *IST3* perturbation observed for artificial introns compared to excised stable introns might suggest that the different 5′- and 3′-end chemistries of the artificial intron conferred greater baseline stability. **e**, The influence of BP–3′SS distance on accumulation of MS2-tagged *UBC4* and *ACT1* introns spliced from *GFP* exons. The diagram shows the 5′-end sequences of the *SAC6* (stable), *UBC4* (stable), *and ACT1* (non-stable) introns, illustrating the absence of a cryptic 5′SS from the *UBC4* and *ACT1* introns. MS2-tagged reporter introns and variants there of that contained altered BP–3′SS distances were expressed in wildtype cells, with those expected to accumulate as stable introns indicated in red. The arrowheads (◂) mark detection of trimmed isoforms of the long BP–3′SS variant, which accumulated in saturated cultures. These isoforms were not labelled by a probe that hybridizes to sequences between the BP and the 3′SS (see Extended Data Fig. 8c), which indicated that they were trimmed at their 3′ termini. Blot is otherwise as described in **c**. *n* = 2 biological replicates. **f**,**g**, Accumulation of T7-driven *UBC4* (**f**) and *ACT1* (**g**) artificial introns; otherwise, as in Fig. 6b. Artificial intron sequences were the same as those expressed from *GFP* mRNA in **e. h**,**i**, Heavy sedimentation of linear artificial introns, even in the absence of debranching activity. In both saturated (**h**) and log-phase (**i**) *dbr1*Δ cultures, linear artificial introns sediment more heavily than excised intron lariats; otherwise, as in Fig. 6e. The migration of the linear artificial-intron isoform to fractions 6–8 matched the migration of stable-intron species that accumulated in wildtype cells (Fig. 6e,f). A band suspected to be a lariat degradation intermediate is indicated (*).

**Extended Data Table 1:**
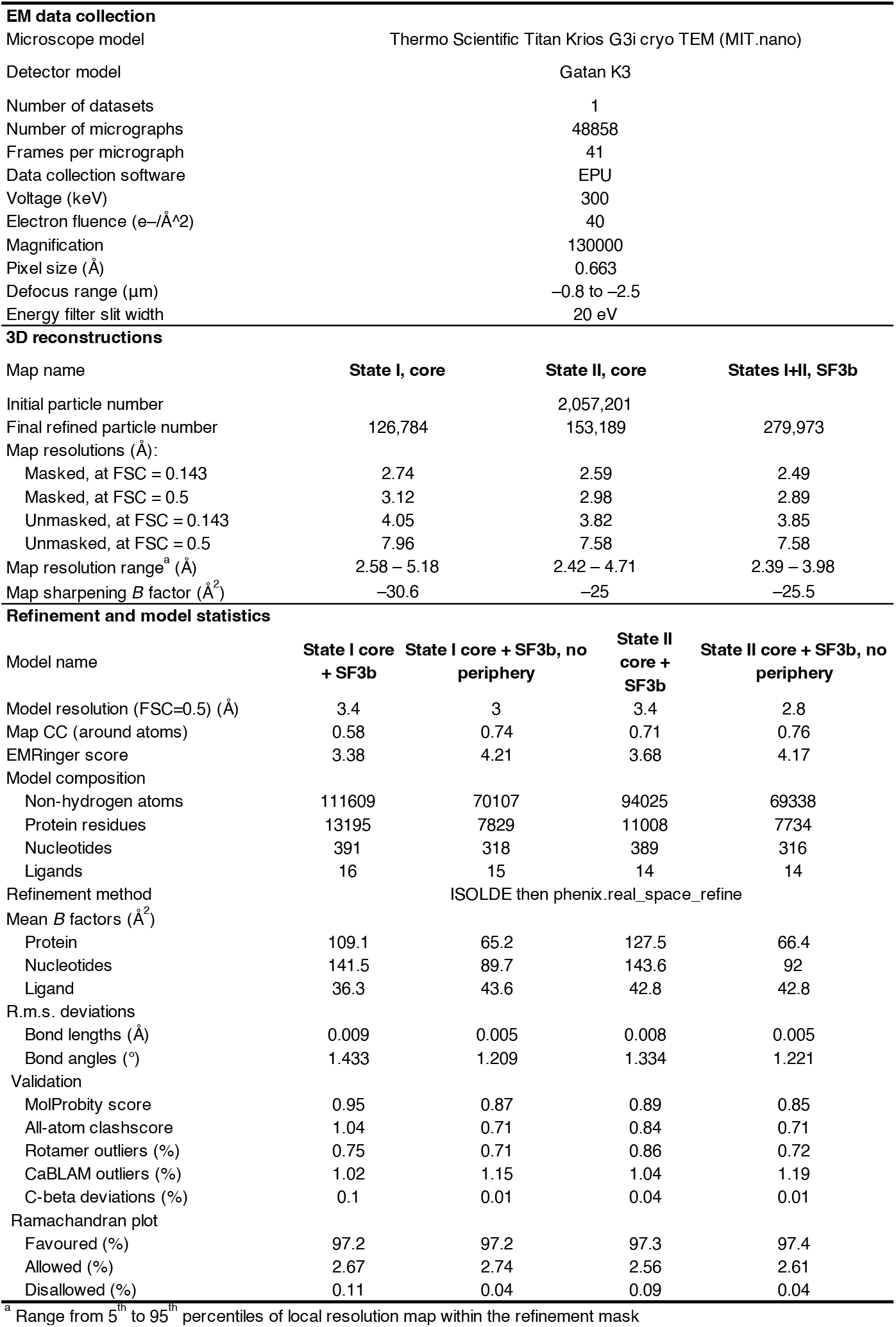
Cryo-EM data collection, refinement and validation statistics

| EM data collection |  |  |  |  |
| --- | --- | --- | --- | --- |
| Microscope model | Thermo Scientific Titan Krios G3i cryo TEM (MIT.nano) |  |  |  |
| Detector model | Gatan K3 |  |  |  |
| Number of datasets | 1 |  |  |  |
| Number of micrographs | 48858 |  |  |  |
| Frames per micrograph | 41 |  |  |  |
| Data collection software | EPU |  |  |  |
| Voltage (keV) | 300 |  |  |  |
| Electron fluence (e-/Å^2) | 40 |  |  |  |
| Magnification | 130000 |  |  |  |
| Pixel size (Å) | 0.663 |  |  |  |
| Defocus range (µm) | −0.8 to −2.5 |  |  |  |
| Energy filter slit width | 20 eV |  |  |  |
| 3D reconstructions |  |  |  |  |
| Map name | State I, core | State II, core | States I+II, SF3b |  |
| Initial particle number |  | 2,057,201 |  |  |
| Final refined particle number | 126,784 | 153,189 | 279,973 |  |
| Map resolutions (Å): |  |  |  |  |
| Masked, at FSC = 0.143 | 2.74 | 2.59 | 2.49 |  |
| Masked, at FSC = 0.5 | 3.12 | 2.98 | 2.89 |  |
| Unmasked, at FSC = 0.143 | 4.05 | 3.82 | 3.85 |  |
| Unmasked, at FSC = 0.5 | 7.96 | 7.58 | 7.58 |  |
| Map resolution range<sup>a</sup> (Å) | 2.58 – 5.18 | 2.42 – 4.71 | 2.39 – 3.98 |  |
| Map sharpening *B* factor (Å<sup>2</sup>) | −30.6 | −25 | −25.5 |  |
| Refinement and model statistics |  |  |  |  |
| Model name | State I core + SF3b | State I core + SF3b, no periphery | State II core + SF3b | State II core + SF3b, no periphery |
| Model resolution (FSC=0.5) (Å) | 3.4 | 3 | 3.4 | 2.8 |
| Map CC (around atoms) | 0.58 | 0.74 | 0.71 | 0.76 |
| EMRinger score | 3.38 | 4.21 | 3.68 | 4.17 |
| Model composition |  |  |  |  |
| Non-hydrogen atoms | 111609 | 70107 | 94025 | 69338 |
| Protein residues | 13195 | 7829 | 11008 | 7734 |
| Nucleotides | 391 | 318 | 389 | 316 |
| Ligands | 16 | 15 | 14 | 14 |
| Refinement method | ISOLDE then phenix.real_space_refine |  |  |  |
| Mean *B* factors (Å<sup>2</sup>) |  |  |  |  |
| Protein | 109.1 | 65.2 | 127.5 | 66.4 |
| Nucleotides | 141.5 | 89.7 | 143.6 | 92 |
| Ligand | 36.3 | 43.6 | 42.8 | 42.8 |
| R.m.s. deviations |  |  |  |  |
| Bond lengths (Å) | 0.009 | 0.005 | 0.008 | 0.005 |
| Bond angles (°) | 1.433 | 1.209 | 1.334 | 1.221 |
| Validation |  |  |  |  |
| MolProbity score | 0.95 | 0.87 | 0.89 | 0.85 |
| All-atom clashscore | 1.04 | 0.71 | 0.84 | 0.71 |
| Rotamer outliers (%) | 0.75 | 0.71 | 0.86 | 0.72 |
| CaBLAM outliers (%) | 1.02 | 1.15 | 1.04 | 1.19 |
| C-beta deviations (%) | 0.1 | 0.01 | 0.04 | 0.01 |
| Ramachandran plot |  |  |  |  |
| Favoured (%) | 97.2 | 97.2 | 97.3 | 97.4 |
| Allowed (%) | 2.67 | 2.74 | 2.56 | 2.61 |
| Disallowed (%) | 0.11 | 0.04 | 0.09 | 0.04 |

<sup>a</sup> Range from 5<sup>th</sup> to 95<sup>th</sup> percentiles of local resolution map within the refinement mask

**Extended Data Table 2:**
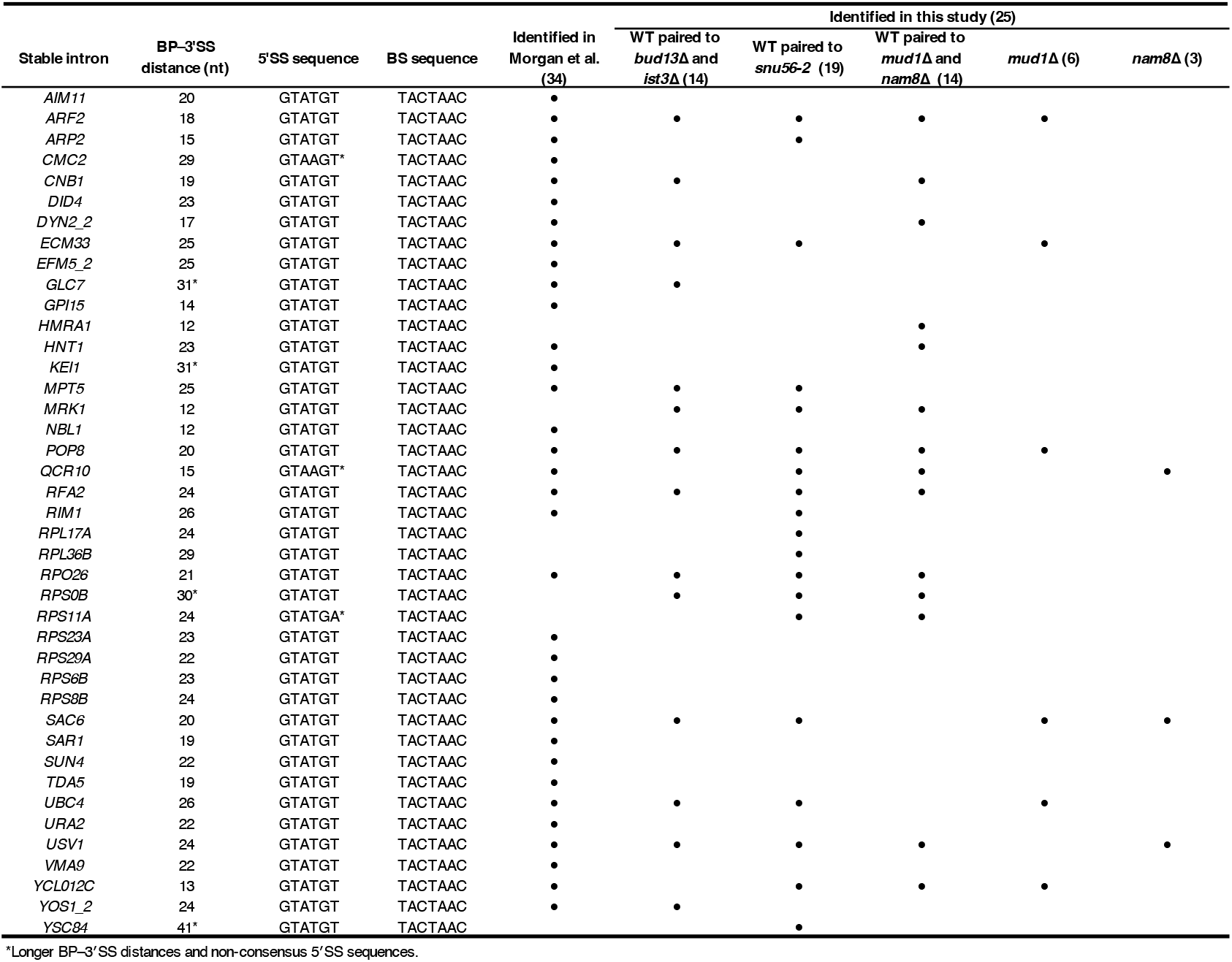
Identities and sequence features of stable introns. Listed are the 34 stable introns originally identified in Morgan et al., (2019), as well as the stable introns annotated in the course of the current study, which brought the total number of annotated stable introns to 41. Indicated for each stable intron is the BP–3 SS distance, the 5 SS sequence, the BS sequence, and the experiment and genetic background in which the intron passed the threshold for de novo annotation from RNA-seq data.

## Supplementary materials

**Supplementary Table 1: S. cerevisiae strains used and generated in this study**. (separate file)

**Supplementary Table 2: Oligonucleotide sequences used in this study**. (separate file)

**Supplementary Table 3: Plasmids used and generated in this study**. (separate file)

**Supplementary Table 4. Mutant intron sequences**. (separate file)

